# WNT6-ACC2-induced accumulation of triacylglycerol rich lipid droplets is exploited by *M. tuberculosis*

**DOI:** 10.1101/2020.06.26.174110

**Authors:** Julius Brandenburg, Sebastian Marwitz, Simone C. Tazoll, Franziska Waldow, Barbara Kalsdorf, Tim Vierbuchen, Thomas Scholzen, Annette Gross, Svenja Goldenbaum, Alexandra Hölscher, Martina Hein, Lara Linnemann, Maja Reimann, Andreas Kispert, Michael Leitges, Jan Rupp, Christoph Lange, Stefan Niemann, Jochen Behrends, Torsten Goldmann, Holger Heine, Ulrich E. Schaible, Christoph Hölscher, Dominik Schwudke, Norbert Reiling

## Abstract

In view of emerging drug-resistant tuberculosis, host directed therapies are urgently needed to improve treatment outcomes with currently available anti-tuberculosis therapies. One option is to interfere with the formation of lipid-laden “foamy” macrophages in the infected host. Here, we provide evidence that WNT6, a member of the evolutionary conserved WNT signaling pathway, promotes foam cell formation by regulating key lipid metabolic genes including acetyl-CoA carboxylase-2 (ACC2) during pulmonary TB. In addition, we demonstrate that *Mycobacterium tuberculosis* (Mtb) facilitates its intracellular growth and dissemination in the host by exploiting the WNT6-ACC2 pathway. Using genetic and pharmacological approaches, we show that lack of functional WNT6 or ACC2 significantly reduces intracellular TAG levels, Mtb growth and necrotic cell death of macrophages. In combination with the anti-TB drug isoniazid, pharmacological inhibition of ACC2 improved anti-mycobacterial treatment *in vitro* and *in vivo*. Therefore, we propose the WNT6-ACC2 signaling pathway as a promising target for a host-directed therapy to reduce intracellular replication of Mtb by modulating neutral lipid metabolism.

## Introduction

Tuberculosis (TB) is the leading cause of death from a single infectious agent^1^. The current increase in the numbers of patients affected by multidrug-resistant (MDR) and rifampicin-resistant (MDR/RR)-TB^2^ severely jeopardizes control of the TB epidemic as envisaged by the WHO “EndTB” strategy^1^. A novel and innovative approach to fight disease without incurring the risk of bacterial resistance development is to target host factors that facilitate *Mycobacterium tuberculosis* (Mtb) replication^3^.

As an intracellular pathogen, Mtb has evolved to reside within the hostile environment of macrophages^4^. These cells serve as the main host cell for Mtb but are also able to restrict infection when appropriately activated. In response to signals such as hypoxia^5^, microbial structures^6^ and Mtb infection^7^, macrophages undergo a substantial metabolic shift away from oxidative metabolism towards glycolysis. Rewiring of cellular metabolism is necessary to mediate macrophage activation^8^, pro-inflammatory polarization^9^ and to control Mtb growth^10,11^. These activating signals, however, also promote the accumulation of neutral lipids in macrophages as fatty acid oxidation is down-regulated^12,13^.

Macrophages with a “foamy”, neutral lipid-rich phenotype are abundantly found in the Mtb-infected human lung and particularly in TB granulomas^14–16^. Moreover, in progressive post-primary TB, infection is restricted to these cells^16,17^. Foamy macrophages accumulate triacylglycerols (TAGs) and cholesterolesters (CEs) in cytoplasmic compartments termed lipid droplets. Cholesterol^18^ and fatty acid^19^ utilization is known to be critical for Mtb growth *in vivo*. Foam cell formation is linked to bacterial persistence, as Mtb is repeatedly found in close proximity to lipid droplets^14^ and utilizes fatty acids derived from host TAGs^20^. Importantly, the presence of foamy macrophages was associated with progressive TB pathology due to a temporal and spatial correlation between the death of foamy macrophages and granuloma evolvement towards tissue necrosis ultimately leading to the release of mycobacteria into the airways^21^. Thus, interfering with foam cell formation during infection may deprive Mtb of essential nutrients within its intracellular niche and restrict bacterial dissemination.

The Wingless/Integrase 1 (WNT) signaling pathway, which is evolutionarily highly conserved in multicellular eukaryotic organisms (metazoa), comprises 19 extracellular WNT ligands in men and mice^22^. WNT signaling regulates basic processes such as proliferation, differentiation and death in virtually all cells including immune cells^23,24^. Previously, we reported that Mtb infection induces expression of WNT6 in macrophages, which acts as an anti-inflammatory feedback regulator dampening responses to mycobacteria^25^. Moreover, we found WNT6 predominantly expressed in a subset of lipid droplet-rich macrophages *in vivo*^25^. In the current study, we demonstrate that WNT6 is a foam cell-promoting factor during Mtb infection. We provide evidence that WNT6-induced acetyl-CoA carboxylase 2 (ACC2) activity in macrophages and mice mediates a metabolic shift away from fatty acid oxidation towards TAG synthesis, which is utilized by Mtb for intracellular replication.

## Results

### WNT6 is expressed in foamy macrophages during pulmonary TB

We have previously reported that WNT6 is expressed in granulomatous infiltrations in the lungs of C57Bl/6 mice experimentally infected with Mtb^25^. To extend this observation to human pulmonary TB, we stained lung tissue samples of three independent TB patients, who have undergone resection of infected lung tissue, with an antibody directed against WNT6 (Figure 1 and S1a-c). WNT6 protein expression was found in cells within nascent granulomas but also in the periphery of necrotizing granulomas (black arrows, Figure 1a). We found WNT6 expression almost exclusively in cells positive for the monocyte/macrophage marker CD68 as revealed by immunofluorescence analyses (Fig. 1c) and immunohistochemical analyses (Fig. S1 a,b). Thus, WNT6 protein expression during Mtb infection in humans is restricted to cells of the myeloid lineage, corroborating previous observations in mice^25^. Of note, WNT6 was prominently expressed in cells with a foam cell morphology (black arrows, Figure 1b). Consistent with that, cells strongly expressing WNT6 (Figure S1d) also showed prominent staining for the lipid droplet scaffolding protein Perilipin2^15^ (PLIN2) (Figure S1e).

**Figure 1:**
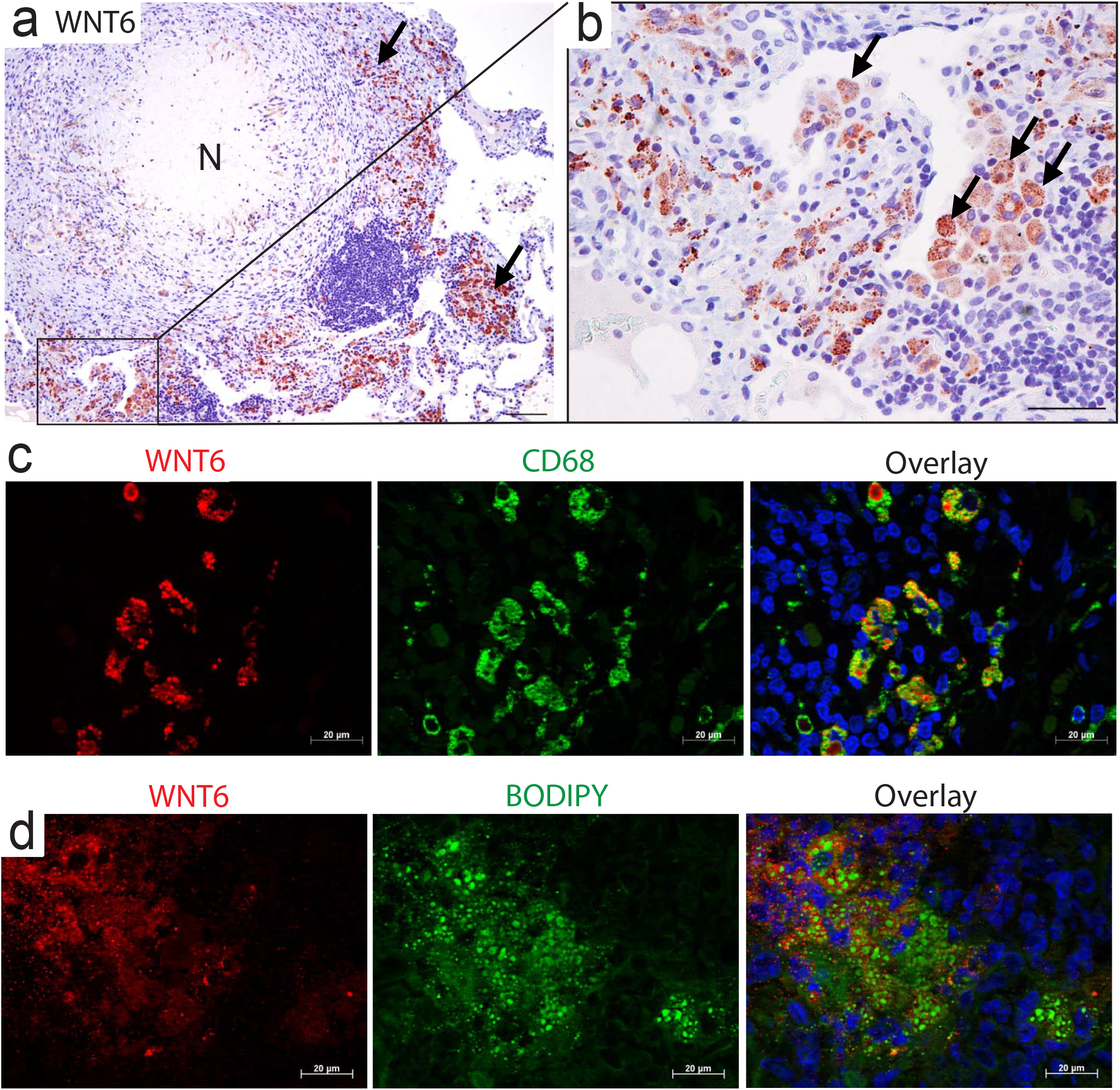
WNT6 is expressed in foamy macrophages during pulmonary TB. **(a-c)** Analyses of formalin-fixed and paraffin-embedded lung tissue derived from a tuberculosis patient. Sections (1 μm) were incubated with an antibody specific for WNT6 (a, b and c, left panel) or the macrophage/monocyte marker CD68^76^ (c, middle panel). Antigens were either visualized with a horseradish peroxidase (HRP)-based detection system using 3-amino-9-ethylcarbazole (AEC) as chromogen (red) (a,b) or by use of specific fluorescence (Cy3 and Cy5)-labeled secondary antibodies (c). Black arrows in (a) indicate areas of WNT6-expressing cells and in (b) cells with a foamy morphology. N, necrosis. Scale bar, (a) 100 μm; (b) 50 μm; (c) 20 μm. **(d)** Immunofluorescence analysis of lung tissue derived from Mtb-infected *IL-13* overexpressing mice (200 CFU; day 63 p.i.). Frozen sections (5 μm) were stained for WNT6 (red) by use of a specific primary antibody and a fluorescence(Cy3)-labeled secondary antibody. Neutral lipids (green) were stained by use of the neutral lipid dye BODIPY 493/503 (10 μg/ml), while nuclei (blue) were counterstained with DAPI. A representative picture of 3 independent observations is depicted.

To further correlate WNT6 expression to the presence of neutral lipids, we analyzed interleukin (IL)-13-overexpressing mice, which develop a human-like pathology upon Mtb infection including centrally necrotizing granulomas with an adjacent zone of foamy macrophages containing numerous lipid droplets (Figure S2a, b and^26^). In these *IL-13* overexpressing mice, an intense WNT6 expression (Figure 1d, left panel, red) was found in areas of prominent neutral lipid accumulation as visualized by staining with the neutral lipid dye BODIPY 493/503^27^ (Figure 1d, middle panel, green and Figure S2c). Together, these findings associate WNT6 to the presence of lipid droplet-rich macrophages in pulmonary TB.

### WNT6 drives accumulation of TAG-rich lipid droplets

We hypothesized that WNT6 expression is functionally linked to the acquisition of a “foamy”, lipid droplet-rich phenotype. To demonstrate this, we analyzed WNT6-overexpressing NIH3T3 cells and visualized neutral lipids by use of BODIPY 493/503. Fluorescence microscopic analysis revealed an enhanced number of neutral lipid-rich structures (Figure 2a, BODIPY, green) in WNT6-overexpressing NIH3T3 cells when compared to control cells, a finding that was independently confirmed by flow cytometry (Figure 2b). Consistent with this, mass spectrometry-based lipid analysis revealed a significantly increased abundance of TAGs in WNT6-overexpressing cells when compared to control cells (Figure 2c). In contrast, the abundance of membrane lipids such as phosphatidylcholines (PCs) remained unchanged (Figure S3a), while the abundance of other neutral lipids species such as cholesterolesters were even decreased (data not shown). To extend these findings to macrophages, Mtb’s main host cells, we next analyzed bone marrow-derived macrophages (BMDMs) from WNT6-competent (*Wnt6*^+/+^) and WNT6-deficient (*Wnt6*^−/−^) mice in the presence of the dietary fatty acid oleic acid conjugated to bovine serum albumin (BSA)^27^. Treatment with oleate-BSA induced lipid droplet formation and enhanced TAG abundance in *Wnt6*^+/+^ BMDMs when compared to control cells as quantified by mass spectrometry (Figure 2e). The presence of lipid droplets and TAG levels were strongly reduced in *Wnt6^−/−^* macrophages when compared to wild-type cells (Figure 2d and 2e). Phosphatidylcholine (PC) levels, which increased upon oleate treatment (PC 36:2), remained comparable between *Wnt6*^+/+^ and *Wnt6*^−/−^ cells (Figure S3b), supporting the notion that WNT6 specifically promotes synthesis of TAG-rich lipid droplets.

**Figure 2:**
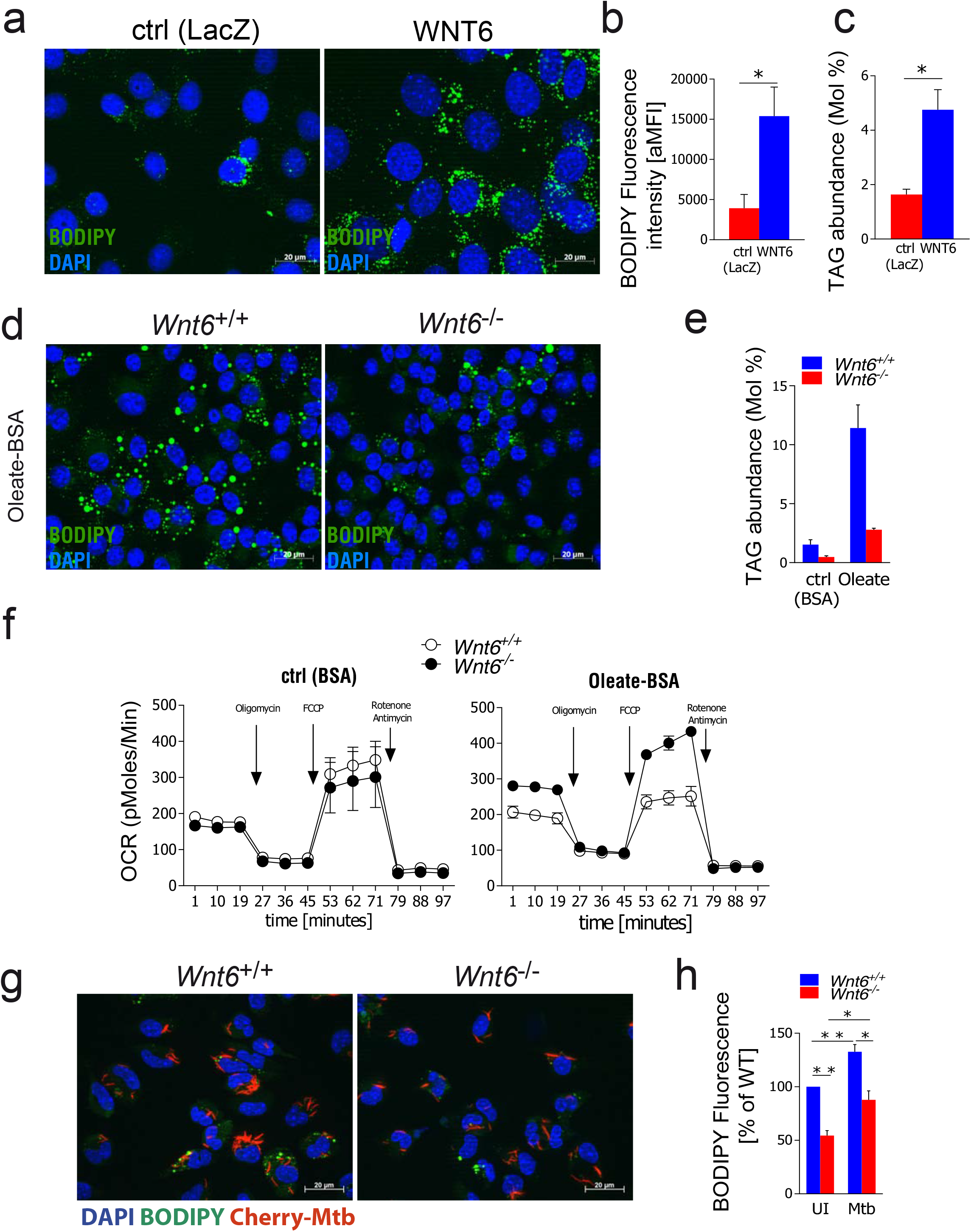
WNT6 drives the accumulation of triacylglycerol (TAG)-rich lipid droplets. **(a)** Visualization of lipid droplets in WNT6-overexpressing or control (LacZ) NIH3T3 cells by fluorescence microscopy; blue, DNA staining (DAPI (1 μg/ml)); green, neutral lipid staining (BODIPY 493/503 (5 μg/ml)); representative staining from 2 independent experiments with similar results is shown. **(b)** Quantification of neutral lipids by flow cytometry (same cells as in (a); arithmetic mean fluorescence intensity (aMFI) of BODIPY signals); n=3. **(c)** Mass spectrometry-based quantification of TAGs (same cells as in (a)); n=3. **(d)** Visualization of lipid droplets in BMDMs from *Wnt6*^+/+^ or *Wnt6*^−/−^ mice incubated for 24 hours in the presence of oleate-BSA (200 μM) (shown is a representative staining from two independent experiments with similar results). **(e)** Mass spectrometry-based quantification of TAGs in BMDMs from *Wnt6*^+/+^ or *Wnt6*^−/−^ mice incubated for 24 h in the presence of BSA (ctrl) or Oleate-BSA (200μM) (n=2). **(f)** Oxygen consumption rate (OCR, pMol per minute) of BMDMs preincubated for 24 h in the presence of BSA (ctrl) or Oleate-BSA (200 μM). Oligomycin (1 μM), FCCP (1.5 μM) and rotenone/antimycin (1 μM); n=2. **(g)** Visualization and **(h)** quantification of neutral lipids in Mtb-infected *Wnt6*^+/+^ and *Wnt6*^−/−^ peritoneal macrophages. After isolation, cells were infected with mCherry-expressing Mtb (MOI 0.1:1) for 24h, stained, and visualized by fluorescence microscopy; red, Mtb; blue, DNA staining (DAPI); green, neutral lipids (BODIPY 493/503). For quantification of neutral lipids (h) over 200 cells per condition were analyzed in 5 independent experiments. Statistical analyses were carried out using One-Way ANOVA with a suitable post-hoc test for multiple comparison. *p≤0.05, **p≤0.01, ***p≤0.001. All data are depicted as mean +/− SEM.

As these results demonstrate that WNT6 regulates macrophage metabolism, we studied the influence of *Wnt6*-deficiency on mitochondrial activity using an extracellular flux analyzer. We observed similar oxygen consumption rates (OCR) between *Wnt6*^+/+^ and *Wnt6*^−/−^ macrophages when cultivating them under control conditions (BSA, Figure 2f, left panel). However, basal as well as maximal respiration measured by the OCR were substantially increased in the absence of WNT6 when treated with oleate-BSA (Figure 2f, right panel). An enhanced oxidative metabolic activity in *Wnt6*-deficient cells upon fatty acid supplementation indicates that WNT6 inhibits mitochondrial fatty acid oxidation and thereby shifts fatty acid metabolism towards neutral lipid synthesis and intracellular storage through the accumulation of lipid droplets.

Next, we assessed whether WNT6 regulates neutral lipid metabolism in *in vivo* differentiated macrophages in the absence or presence of Mtb. We isolated peritoneal macrophages from *Wnt6*^+/+^ and *Wnt6*^−/−^ mice, which were infected with mCherry-expressing Mtb for 24h and analyzed by fluorescence microscopy (Figure 2g,h). Neutral lipid levels as determined by BODIPY 493/503 staining were significantly lower in uninfected cells from *Wnt6*^−/−^ mice compared to *Wnt6*^+/+^ mice (Figure 2h), suggesting that disrupted WNT6 signaling affects neutral lipid levels under homeostatic conditions. Mtb infection - independently of the presence of WNT6 - enhances the amounts of intracellular neutral lipids by ~30% when compared to uninfected cells (Figure 2h), which is consistent with data from a recent study^28^. A significant reduction of BODIPY fluorescence by ~45% was observed in both, uninfected and Mtb-infected peritoneal macrophages from *Wnt6*^−/−^ mice when compared to respective *Wnt6*^+/+^ cells (Figure 2h). Our data show that WNT6-dependent and WNT6-independent pathways contribute to the accumulation of lipid droplets in *in vivo* differentiated macrophages. Taken together, our findings demonstrate that WNT6 drives the accumulation of TAG-rich lipids droplets in the absence and presence of Mtb.

### WNT6 induces expression of lipid metabolic enzymes critical for TAG synthesis and lipid droplet accumulation

To identify the cellular processes that are altered in the absence of *Wnt6,* we conducted a microarray-based gene expression analysis comparing Mtb-infected *Wnt6*^+/+^ and *Wnt6*^−/−^ macrophages. By performing a gene set enrichment analysis (GSEA)^29^ utilizing gene sets from the peer-reviewed Reactome pathway database, we identified “Metabolism of Lipids and Lipoproteins” on rank 4 (FDR q-value, 2.58e^−13^) under the top 10 of enriched gene sets (Figure S4a) along with other expected sets of genes such as “Immune system”, “Cell cycle” and “Development Biology”, which corroborate previous data^25^. In-depth analysis revealed that genes encoding key factors involved in fatty acid uptake (*Cd36,* cluster of differentiation 36)^30^, activation (*Acsl5*, long-chain-fatty-acid-CoA ligase 5)^31^, and mitochondrial oxidation (*Acad8*, acyl-CoA dehydrogenase family member 8,) are significantly up-regulated in *Wnt6*^−/−^ cells (Figure 3a). Of note, *Wnt6*^−/−^ cells showed a strong up-regulation of *Cpt1b*, a gene encoding for an isoform of CPT1, the rate-limiting enzyme in mitochondrial beta-oxidation^32^ (Figure 3a). Consistent with these observations, *Wnt6*^−/−^ cells exhibited decreased mRNA expression of genes associated with fatty acid synthesis (*Acot1*, acyl-CoA thioesterase 1^33^; *Fads6*, fatty acid desaturase 6; *Elovl2*, elongation of very long chain fatty acids 2)^34^) and storage of fatty acids or other lipids (*Bdh1*, 3-hydroxybutyrate dehydrogenase 1) (Figure 3b). Moreover, expression of the key enzyme in TAG synthesis acyl-CoA:diacylglycerol acyltransferase (*Dgat2*)^35^ and the lipid droplet scaffolding protein perilipin3 (*Plin3*)^36^ were down-regulated when compared to *Wnt6*^+/+^ cells. Microarray data also identified a strongly reduced expression of acetyl-CoA carboxylase-2 (*Acacb*, ACC2) (Figure 3b), which acts as a key regulator of fatty acid oxidation through CPT1 inhibition^37,38^ thereby promoting cellular lipid storage^39^.

**Figure 3:**
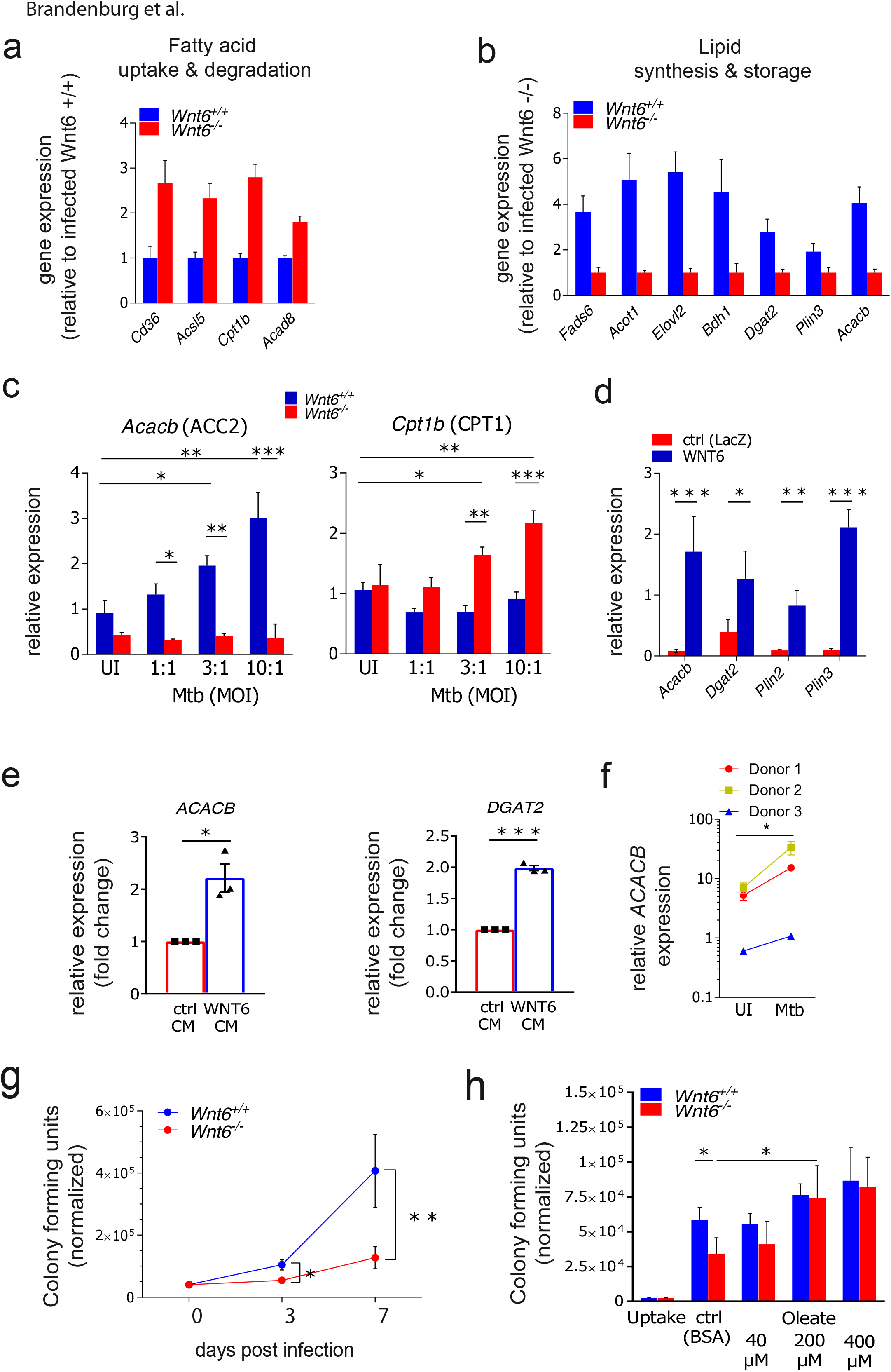
WNT6-mediated changes in host lipid metabolism promote Mtb growth in macrophages. **(a,b)** Microarray-based gene expression analysis of *Wnt6*^+/+^ and *Wnt6*^−/−^ BMDMs infected for 24 hours with Mtb H37Rv (MOI 3:1). Fold expression of statistically significantly regulated genes associated with fatty acid uptake and degradation **(a)** or lipid synthesis and storage **(b)** are depicted; n=3. **(c)** qRT-PCR based gene expression analysis of *Wnt6*^+/+^ and *Wnt6*^−/−^ BMDMs infected for 24 h with various doses (MOIs) of Mtb H37Rv; n=3. **(d)** qRT-PCR based gene expression analysis of WNT6-overexpressing (WNT6) or control (ctrl (LacZ)) NIH3T3 cells; n=3. **(e)** qRT-PCR based gene expression analysis of hMDMs treated with WNT6 conditioned medium (WNT6 CM) or control conditioned medium (ctrl) CM for 24 hours. Fold change relative to control (ctrl CM) is shown. For statistical comparison, raw data were used. Data from 3 independent experiments using cells from different donors are shown; n=3. **(f)** qRT-PCR based gene expression analysis of *ACACB* (ACC2) mRNA expression in Mtb-infected hMDMs at day 7 p.i‥ Cells were infected with Mtb H37Rv (MOI) 1:1), washed (4 h p.i.) and incubated for 7 days; n=3. **(g)** CFU analysis of Mtb-infected (MOI 1:1) *Wnt6*^+/+^ or *Wnt6*^−/−^ BMDMs at day 0 (4 h), 3 and 7 p.i. **(h)** CFU analysis of Mtb-infected (MOI 0.1:1) *Wnt6*^+/+^ or *Wnt6*^−/−^ BMDMs at day 7 p.i. after incubation of cells various concentrations of oleic acid (oleate-BSA). Bacterial growth was related to the number of macrophages (normalized CFU) at the individual timepoint/condition (given as CFU per 100.000 cells). Shown is the mean +/− SEM of a total of 3 **(g)** or 4 **(h)** independent experiments. Statistical analyses were carried out using One-Way ANOVA with a suitable post-hoc test for multiple comparison except microarray-based gene expression analysis **(c,d),** which was conducted as described in *Material and Methods*. *p≤0.05, **p≤0.01, ***p≤0.001. All data are depicted as mean +/− SEM.

Validation by qRT-PCR confirmed that there is indeed an inverse correlation between the expression of *Acacb* and *Cpt1b* depending on the presence of WNT6 (Figure 3c). *Acacb* expression levels were increased upon Mtb infection (24 hours p.i.) in *Wnt6*^+/+^ cells in a MOI dependent manner, while mRNA levels remained at baseline in *Wnt6*^−/−^ macrophages at all MOIs tested. The opposite was observed regarding *Cpt1b* mRNA levels, which were significantly increased in *Wnt6*^−/−^ macrophages upon infection when compared to respective *Wnt6*^+/+^ cells (Figure 3c). *Plin3* mRNA levels were up-regulated upon infection with Mtb in a dose-dependent manner, while *Plin3* mRNA levels remained largely unchanged in the absence of *Wnt6* (Figure S4b). Consistent with these findings, a strongly enhanced expression of *Acacb*, *Dgat2, Plin2* and *Plin3* was observed in WNT6-overexpressing (WNT6) NIH3T3 cells when compared to control (ctrl (LacZ) cells (Figure 3d), providing further evidence that WNT6 drives the expression of key metabolic factors associated with TAG synthesis (ACC2^37–39^ and DGAT2^35^) and lipid droplet biogenesis (PLIN2^15^ and PLIN3^36^).

To investigate whether WNT6 regulates ACC2 expression also in human cells, we analyzed WNT6 conditioned media (WNT6 CM) or control conditioned media (ctrl CM) treated human monocyte-derived macrophages (hMDMs) by qRT-PCR (Figure 3e). After 24h incubation, exogenous WNT6 induced the mRNA expression of ACC2 (*ACACB*) and DGAT2 (*DGAT2)* by ~2fold, which is consistent with results from murine cells (compare to Figure 3b and 3c). Next, we investigated whether Mtb induces ACC2 expression in hMDMs. qRT-PCR analysis of macrophages after 7 days of culture revealed that *ACACB* mRNA expression levels vary substantially between individual human donors (see uninfected (UI), Figure 3f). Upon Mtb infection, *ACACB* mRNA levels were not altered at the 24h timepoint (data not shown), whereas at days 4 (data not shown) and 7 post infection (Figure 3f), a statistically significant increase of *ACACB* mRNA levels was observed albeit in a donor-dependent manner (fold increase between 1.7-4.8). Together, these data show that WNT6 drives expression of key lipid metabolic enzymes, including ACC2, in both murine and human cells.

### WNT6-mediated changes in host lipid metabolism promote Mtb growth in macrophages

To assess whether WNT6-mediated changes in host lipid metabolism affect Mtb’s ability to replicate intracellularly, we addressed Mtb growth in *Wnt6*^+/+−^ and *Wnt6*^−/−^ macrophages. The number of Mtb bacteria 4h p.i. was comparable between both cell types independent of the dose of infection (Figure S4c). However, intracellular bacterial loads were significantly reduced in cells lacking *Wnt6* (Figure 3g and Figure S4d) at day 3 p.i. (MOI 1, Figure 3g), showing approximately 50% reduced bacterial numbers in *Wnt6*^−/−^ cells when compared to *Wnt6*^+/+^ macrophages. Bacterial loads in *Wnt6*^+/+^ cells further increased until day 7 post infection, while CFUs in *Wnt6*^−/−^ cells remained at a rather low level (Figure 3g). This amounts to a CFU reduction of approximately 70% in the absence of *Wnt6*. At both time points analyzed, the quantification of nitrite in cell culture supernatants of *Wnt6*-competent and *Wnt6*-deficient macrophages revealed similar production of nitric oxide, a well-established tuberculostatic host factor (Figure S4e). Moreover, acidification rates of Mtb-containing compartments were similar between *Wnt6*^+/+^ and *Wnt6*^−/−^ macrophages as determined by fluorescence microscopy analyses of the intracellular localization of GFP-Mtb (green) and LysoTracker Dye (red) (Figure S4f). These findings suggest that WNT6 promotes Mtb growth without affecting bacterial uptake, phagosome acidification, and nitric oxide production of infected macrophages.

In order to test whether a reduced availability of lipid substrates is the cause for the impaired growth of Mtb in *Wnt6*^−/−^ cells, we supplemented macrophage cultures with various concentrations of oleate-BSA and determined CFU development on day 7 p.i. (Figure 3h). CFU levels in ctrl (BSA)-treated cultures were reduced by 42% in *Wnt6*^−/−^ macrophages when compared to *Wnt6*^+/+^ cells. Addition of 200 μM oleate-BSA to *Wnt6*^−/−^ macrophages led to significantly enhanced CFU numbers, which were similar to those in *Wnt6*^+/+^ BMDMs. Higher oleate-BSA concentrations (400 μM) also led to a comparable bacterial burden in *Wnt6*^+/+^ and *Wnt6*^−/−^ cells. These data strongly suggest that WNT6-dependent changes in the cellular availability of lipids promote Mtb growth in macrophages.

### ACC2 activity promotes bacterial growth in macrophages

To assess whether the identified WNT6 target enzyme ACC2 promotes Mtb growth in macrophages, we generated functional protein knockouts of both isoforms, ACC1 and 2, by CRISPR/Cas9 mediated genome-editing in the human macrophage-like cell line BLaER1^40,41^. Mtb growth analyses revealed that deficiency of ACC2 but not of ACC1 significantly reduces Mtb CFUs at day 3 p.i. (by ~58%) when compared to wild-type (WT) cells (Figure 4a). To substantiate this finding, we treated primary human macrophages (hMDMs) with three structurally different pharmacological ACC2 inhibitors. All tested compounds reduced Mtb growth dose-dependently when compared to solvent control, albeit with varying efficacy (ranging from ~29-84% growth reduction, Figure 4b-e). Of note, the inhibitors tested did not exert toxic effects on human macrophages (Figure S5a and data not shown). The inhibitor concentrations used also did not inhibit Mtb growth in liquid culture as indicated by comparable fluorescence signals between ctrl (solvent) or ACC2 inhibitor treated mCherry-expressing Mtb bacteria (see Figure S5b). Moreover, we did not observe a direct effect of ACC2 inhibition on the immediate inflammatory response of hMDMs to Mtb as determined by measurement of TNFα release at day 1 (Figure S5c), 4 or 7 p.i. (data not shown). Taken together, our findings show that genetic and pharmacologic targeting of ACC2 activity restricts Mtb growth within human macrophages without reducing viability or the pro-inflammatory response of these cells. In order to test whether a reduced availability of lipid substrates is also the cause for the impaired growth in cells lacking active ACC2, we determined intracellular growth upon addition of fatty acids in the absence and presence of ACC2 inhibitors. This analysis revealed that both oleate as well as palmitate promote Mtb growth when added to hMDM cultures (Figure S5d). Exogenously added fatty acids – depending on the efficacy of the inhibitor and fatty acid used - can restore Mtb growth in ACC2 inhibititor treated cells (Figure 4f), suggesting a functional link between ACC2 dependent availability of cellular lipids and intracellular Mtb replication. Targeting ACC2 as a host-metabolic enzyme could complement pathogen-directed antibiotic treatments of TB. Thus, we tested the growth-inhibiting effect of ACC2 inhibition on Mtb growth in primary human macrophages in combination with the first line anti-TB drug isoniazid (INH), which was applied at suboptimal concentration (0.03 μg/ml). In the same set of experiments already shown before (Figure 4b), ACC2 inhibitor 1 or INH alone lead to a growth reduction of ~89% and ~79%, respectively, when compared to solvent control (Figure 4c). Treating cells with a combination of both resulted in a growth reduction of ~96% (Figure 4c), revealing a nearly additive effect of these drugs.

**Figure 4:**
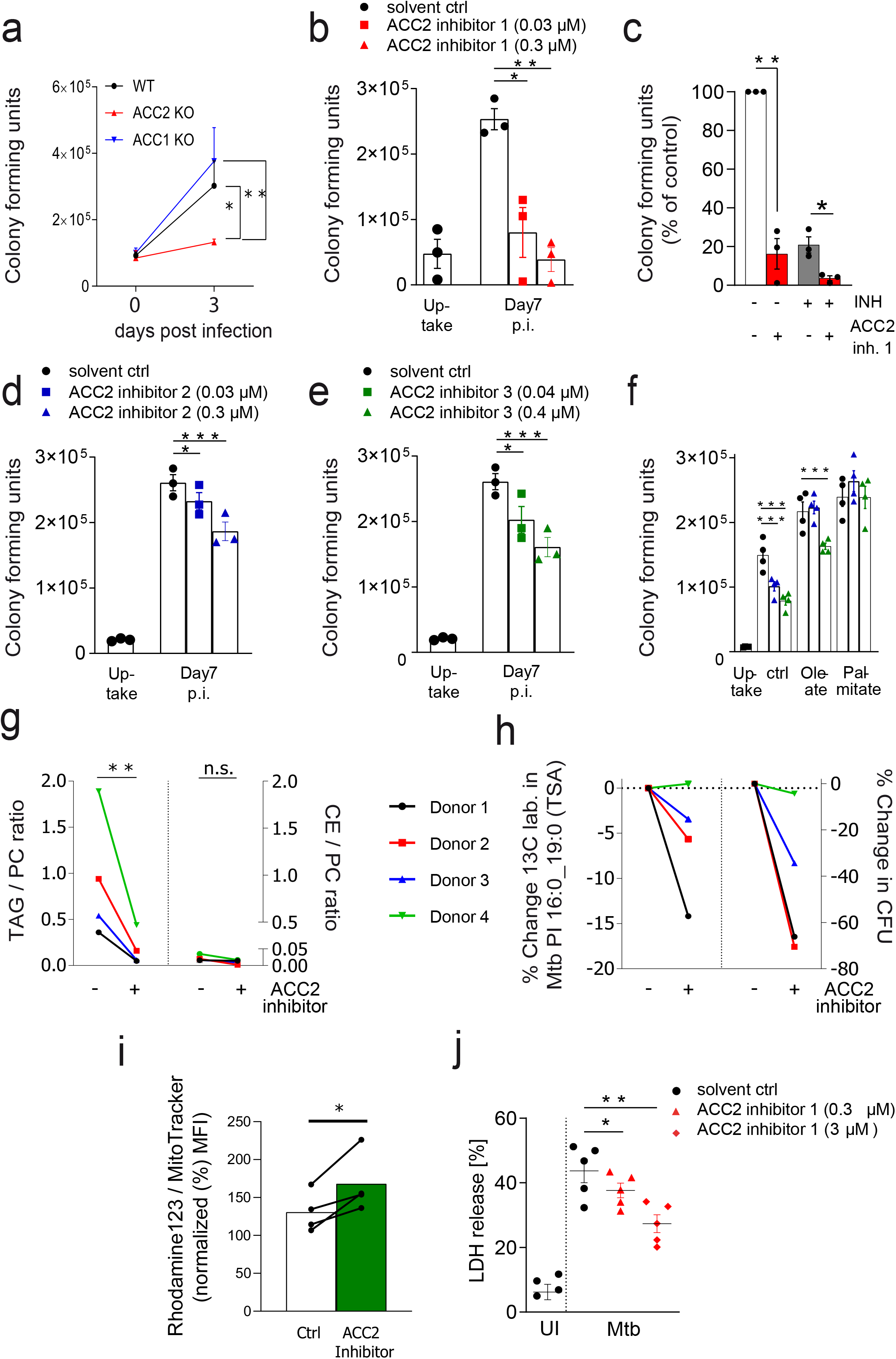
ACC2 activity promotes bacterial growth in macrophages by promoting TAG accumulation and necrotic cell death. **(a)** CFU analysis of Mtb-infected (MOI 0.5:1) wild-type (WT), ACC1 KO and ACC2 KO human macrophage-like cells (BLaER1 macrophages) at day 3 p.i‥ n=3. **(b,d,e)** CFU analysis of Mtb-infected (MOI 1:1) hMDMs treated with pharmacological ACC2 inhibitors at day 7 p.i‥ Uptake was determined 4h p.i‥ After washing, cells were incubated in the absence (solvent ctrl, DMSO) or presence of different ACC2 inhibitors with the concentrations indicated; n=3. **(c)** In the same set of experiments shown in (b), additionally to the treatment with ACC2 inhibitor 1 alone (300nM), cells were also treated with isoniazid (INH (0.03 μg/ml)) or as a combination of ACC2 inhibitor plus INH. Shown is the relative reduction of CFU (%). **(f)** CFU analysis of Mtb-infected (MOI 0.5:1) hMDMs treated with ACC2 inhibitors in the presence of exogenous fatty acids at day 7 p.i‥ Uptake was determined 4h p.i‥ After washing, cells were incubated with oleate-BSA or palmitate-BSA (both 400μM) in the absence or presence of ACC2 inhibitors 2 (blue triangles, 300 nM) or ACC2 inhibitor 3 (green triangles, 400 nM); n=4. **(g,h)** hMDMs were pulsed with isotopically labelled ^13^C-Oleate-BSA prior to infection with Mtb (MOI 1:1) and subsequently incubated in the absence (solvent, ctrl) and presence of ACC2 inhibitor 3 (400 nM) for 7 days. Mass spectrometry-based analyses show (g) ratios of TAG (left panel) and CE (right panel) normalized to PC and (h, left panel) the change (%) in isotope labeling (^13^C18) in the Mtb-specific membrane lipid PI 16:0_19:0 tuberculostearic acid (TSA) from the same sample. In parallel, cells were subjected to CFU analysis, revealing the % of CFU reduction (h, right panel) from each individual experiment (donor); n=4. **(i)** Flow cytometry-based quantification of - Rhodamine 123 signals (relative to MitoTracker Deep Red signals (both aMFI) x100) in Mtb-infected (MOI 0.1:1) and ACC2 inhibitor 3 (400nM) treated hMDMs at day 3 p.i.; n=4. **(j)** Quantification of Lactate Dehydrogenase Release (LDH) from hMDM cultures at day 7 p.i.; n=5. Cells were equally infected as described for (b). UI, uninfected; TAG, Triacylglycerols; PC, Phosphatidylcholines; CE, Cholesterolester. Statistical analyses were carried out using One-Way ANOVA with a suitable post-hoc test for multiple comparison; *p≤0.05, **p≤0.01, ***p≤0.001. All data are depicted as mean +/− SEM.

### ACC inhibition lowers TAG levels in infected macrophages and utilization of host cell fatty acids by Mtb

In order to address whether ACC2 inhibition affects Mtb’s ability to utilize host cell lipids from macrophages, we pulsed human macrophages with ^13^C-labelled oleate prior to infection with Mtb. Mass spectrometric analyses demonstrated that the labeled oleate was effectively incorporated into TAG, CE and PC species of the host cell (Figure S6a and Supplementary Table I). A comparative analysis of Mtb infected cells at d7 p.i. showed that ACC2 inhibitor treatment – compared to solvent control – led to an overall reduction of TAG/PC ratios in macrophages (Figure 4g, left panel and Supplementary Table I), whereas CE/PC ratios - present in a drastically lower abundance - remained unchanged (Figure 4g, right panel). This shows that primarily TAGs are affected by inhibition of ACC2 enzyme activity. To trace the fate of ^13^C-labelled oleic acid in Mtb, we monitored the incorporation of the labelled substrate into tuberculostearic acid (TSA, C19:0), a characteristic fatty acid of acid-fast bacteria of the order Actinomycetales^42 43^. In an independent study, we have established the detection and quantification of TSA in a highly abundant cell membrane phosphatidylinositol of Mtb (PI 16:0_19:0 (TSA)) (preprint: Heyckendorf et al. Biorxiv, 2020). In the current study, we found that ^13^C-labelling in Mtb PI 16:0_19:0 (TSA) (Figure S6b, lower panel) is reduced in samples from 3 out of 4 donors upon ACC2 inhibition (Figure 4h, left panel), which showed low TAG/PC ratios of 0.04, 0.11 and 0.04 (Figure 4g). This correlated with the magnitude of Mtb growth reduction (Figure 4h, right panel). Both, ^13^C-labeling in Mtb and CFUs remained almost unchanged in samples from donor 4, which showed an up to 8fold increased TAG/PC ratio of 0.34 (Figure 4g), when compared to donor 1,2 and 3. Collectively, these data show that Mtb metabolizes host cell fatty acids, the metabolization of which is reduced when host ACC2 is inhibited. These findings suggests that intracellular replication of Mtb requires sufficient access to TAG-derived lipid nutrients.

### ACC2 inhibition enhances mitochondrial activity and limits Mtb-induced necrotic cell death of macrophages

ACC2 activity is known to impair mitochondrial fatty acid oxidation through CPT1 inhibition^37–39^. Consistent with that, mitochondria were metabolically more active in ACC2 KO BlaER1 macrophages (Figure S5e) as well as in ACC2-inhibitor treated cells as indicated by increased relative fluorescence signals of the membrane potential sensitive fluorochrome rhodamine 123^44,45^ in Mtb-infected, ACC2 inhibitor-treated primary human macrophages when compared to control cells (day 3 p.i., Figure 4i). Based on these data and the previous observation that enhanced fatty acid oxidation upon ACC inhibition protects cells against lipotoxicity^46^, we monitored the viability of Mtb-infected macrophages in the absence or presence of ACC2 inhibitors. Indeed, we microscopically observed that ACC2-inhibitor treatment prolonged the survival of Mtb infected macrophages (Figure S5f). This prompted us to analyze necrotic cell death by measuring release of lactate dehydrogenase (LDH) as marker of cell membrane disruption^47^. Macrophages infected with Mtb showed a marked increase in LDH release during the course of infection in a time dependent manner (data not shown) with a ~43% maximum release at day 7 p.i. (Figure 4j). Strikingly, when treating infected cells with ACC2 inhibitor we observed a statistically significant reduction of LDH release in a dose-dependent manner when compared to solvent control (up to ~50% reduction of LDH release, Figure 4j). Taken together our findings suggest that WNT6-driven ACC2 activity is instrumental in promoting TAG accumulation and contributes to lipotoxicity-induced necrotic cell death in macrophages, both of which contribute to Mtb replication and dissemination in the infected host^48–50^.

### ACC2 inhibition improves anti-mycobacterial treatment *in vivo*

We then explored the presence of ACC2 *in vivo* and analyzed ACC2 expression in lung tissue sections of a TB patient. A strong ACC2 signal was found in the periphery of human necrotizing granulomas coinciding with the presence of CD68^+^ cells (see boxes in Figure 5a and Figure S7a), suggesting that ACC2 plays a role during active TB in humans. Finally, we investigated the functional role of ACC2 *in vivo* employing an experimental murine model of TB infection. To our surprise, immunohistochemical stainings did not reveal a prominent ACC expression in the lungs of C57Bl/6 mice, even when infected with a high dose of Mtb (Figure 5c). In contrast, numerous ACC positive cells were easily detectable in low dose infected 129/Sv mice (Figure 5d), which are known to develop a TB susceptible phenotype resembling primary progressive TB disease in humans^51^.

**Figure 5:**
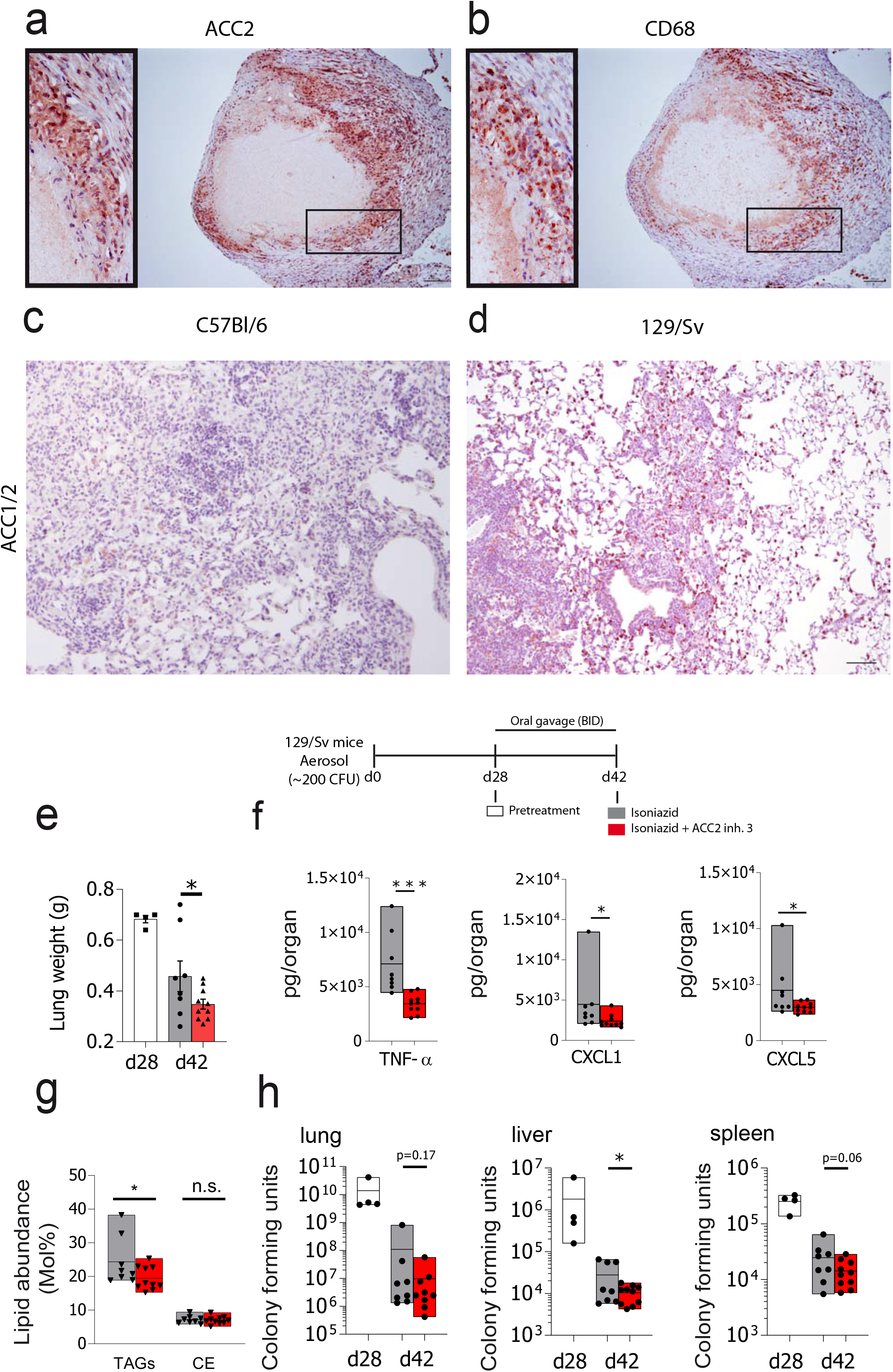
ACC2 expression affects disease development in pulmonary TB. **(a,b)** Immunohistochemical analyses of formalin-fixed and paraffin-embedded lung tissue derived from a tuberculosis patient. Consecutive sections (1μm) were incubated with antibodies specific for ACC2 (a) or the macrophage/monocyte marker CD68 (b). Antigens were visualized with a Horseradish peroxidase (HRP)-based detection system using AEC as chromogen (red). Scale bar, 100 μm **(c,d)** Immunohistochemical analyses of formalin-fixed and paraffin-embedded lung tissue of Mtb-infected C57Bl/6 (~1000 CFU, d42 p.i.) or 129/Sv mice (~200 CFU, d28 p.i.). Sections (2 μm) were incubated with antibodies specific for ACC 1/2 and antigens visualized with a Horseradish peroxidase (HRP)-based detection system using AEC as chromogen (red). **(e-h)** *In vivo* efficacy of ACC2 inhibitor treatment when combined with the first-line anti-TB drug INH. After 28 days of infection with Mtb H37Rv (~200CFU), 129/Sv mice were either left untreated (pretreatment, d28 p.i., n=4, white bars) or were treated for 14 days with INH alone (10 mg/ per kg bodyweight (BW), n=8, grey bars) or with ACC2 inhibitor 3 (ND-646, 25 mg/kg BW) plus INH (n=10, red bars). Lung weights (e), lung cytokine and chemokine levels (f), TAG and CE abundance in the lung (g), as well as mycobacterial loads in lung, liver and spleen were determined. Statistical analyses were carried out using an one-tailed, unpaired Student’s t-test; *p≤0.05, **p≤0.01, ***p≤0.001; n.s.= not significant. Data are depicted as Min-Max bar with line at mean.

To evaluate the efficacy of ACC2 inhibition *in vivo*, 129/Sv mice infected with Mtb for 28 days were subjected to a short-term, low-dose treatment (25mg/kg BW) with ACC2 inhibitor 3. This compound has been successfully tested in a preclinical mouse model of lung cancer^52^. Seven days after beginning of treatment with the ACC2 inhibitor, no substantial changes with regard to the inflammatory response and the bacterial burden was observed, when homogenates of lung, liver and spleen of ACC2 inhibitor treated mice were compared to those of vehicle control treated mice (Figure S7b and data not shown).

Our *in vitro* data reveal a strong Mtb growth reducing effect when INH and ACC2 inhibitors were added to macrophage cultures simultaneously (Figure 4c). Since targeting of host ACC2 would always be an adjunct to standard TB therapy, we combined ACC2 inhibitor with INH *in vivo*. A dose of 10 mg/kg bodyweight (BW) was chosen, as this is comparable to drug plasma concentrations observed in humans rapidly metabolizing INH (rapid acetylator phenotype)^53,54^. Two weeks after starting treatment, the concomitant administration of INH and the ACC2 inhibitor significantly reduced lung weights of infected mice when compared to INH treated animals by 25% (Figure 5e), which is indicative for a reduced presence of inflammatory cells in the lungs of these mice. In line with these results the combination of INH plus ACC2 inhibitor significantly reduced the production of the major pro-inflammatory cytokine TNFα as well as the neutrophil chemoattractants CXCL1 and CXCL5 (Figure 5f), when compared to mice treated with INH alone (day 42 p.i.). The expression levels of these chemokines correlate with bacterial loads and disease severity in susceptible Mtb-infected mice and have been associated with lung and granuloma necrosis^55^.

Moreover, mice treated with INH plus ACC2 inhibitor showed significantly reduced TAG abundance in the lungs, but no changes in the amounts of cholesterol esters (Figure 5g), when compared to mice treated with INH alone, demonstrating that ACC2 inhibition affects TAG levels in the infected mouse lung. With regard to the impact on the bacterial burden, we observed that two weeks after starting treatment, INH and the ACC2 inhibitor together substantially reduced mean Mtb CFUs in lung (10fold), liver (2.7fold) and spleen (1.7fold) when compared to INH alone (Figure 5h). This effect reached statistical significance in the liver, but not in the lung (p=0.17) and spleen (p=0.06). Taken together, our data suggest that even a late and limited adjunct treatment with a pharmacological ACC2 inhibitor has an impact on the course of experimental Mtb infection *in vivo*.

## Discussion

Foamy macrophages are key players in TB as they provide a nutrient-rich reservoir for mycobacterial replication and contribute to tissue pathology^21^. However, the detailed mechanisms of how Mtb infection induces the development of these lipid-laden cells are still unclear. Our findings show that the WNT ligand WNT6 acts as a foamy macrophage-promoting factor in pulmonary TB by inducing acetyl-CoA carboxylase-2 (ACC2) (summarized in Figure 6). We found prominent WNT6 expression in cells showing characteristics of foamy macrophages in pulmonary granulomas from TB patients, as well as in mice, which develop human-like granuloma necrosis upon Mtb-infection^26^. In terms of function, our study reveals that the WNT6-ACC2 metabolic axis drives the accumulation of lipid droplets containing high levels of TAGs. We demonstrate that inhibiting ACC2 reduces TAG concentrations in macrophages *in vitro* as well as in the lungs of Mtb-infected mice *in vivo*. Moreover, our findings that the lack of *Wnt6* or ACC2 reduce intracellular Mtb replication in macrophages strongly suggest that WNT6-ACC2 induced changes in neutral lipid metabolism affect disease progression. Indeed, pharmacological ACC2 inhibitors, when combined with the first line drug isoniazid, improved anti-mycobacterial treatment in infected macrophages and mice. Together, our findings show that WNT6-ACC2 dependent metabolic changes leading to accumulation of TAGs in macrophages are exploited by the pathogen to facilitate its intracellular replication.

**Figure 6:**
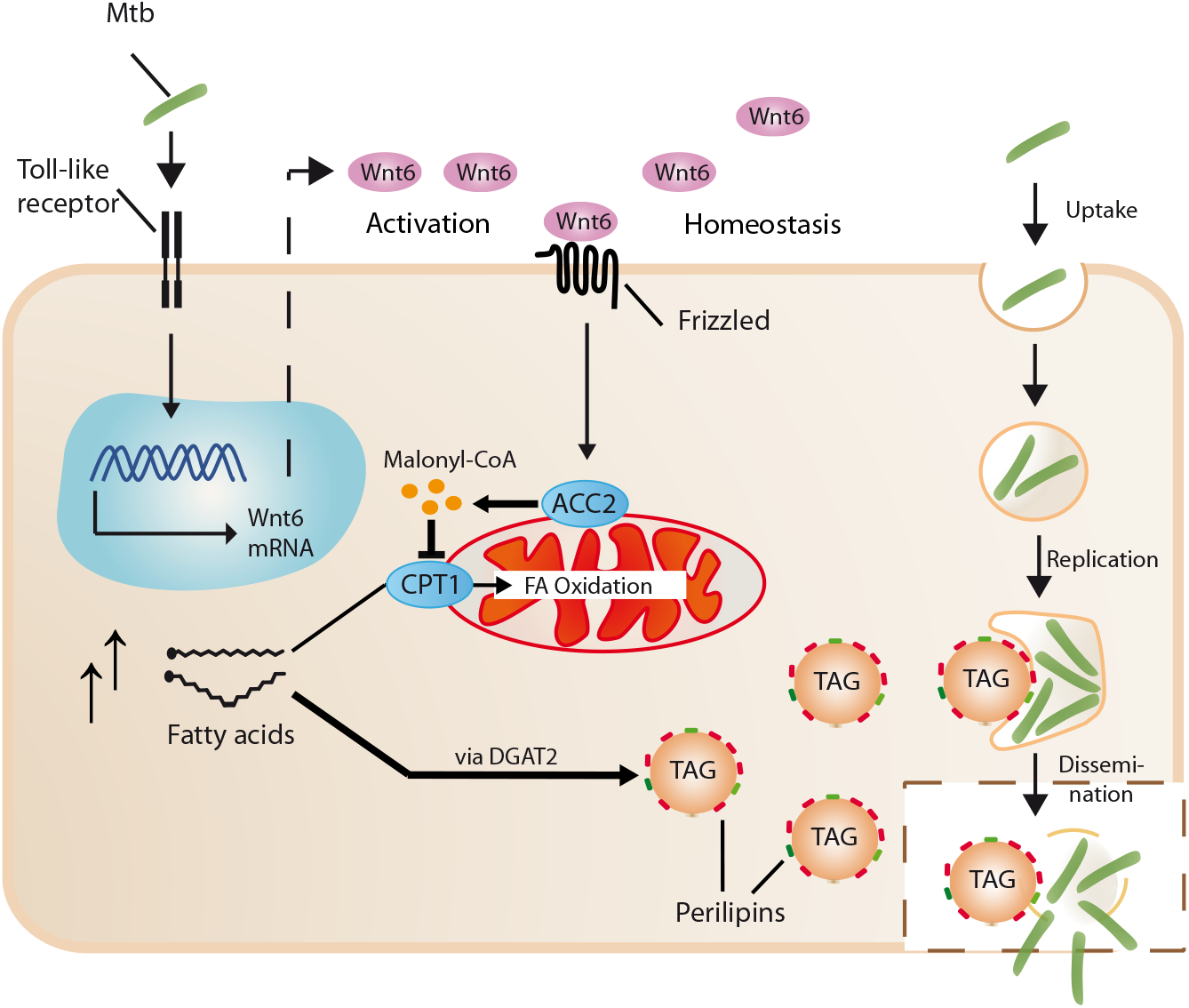
WNT6-ACC2-induced accumulation of triacylglycerol rich lipid droplets is exploited by *M. tuberculosis*. Homeostatic-or activation (TLR2/4)-dependent WNT6-signaling via Frizzled receptors induces the expression of various key lipid metabolic genes including acyl-CoA:diacylglycerol acyltransferase (DGAT2) and Acetyl-CoA Carboxylase-2 (ACC2). ACC2 is known to generate Malonyl-CoA, which inhibits carnitine palmitoyltransferase 1 (CPT1)-dependent import of fatty acids into mitochondria thereby reducing cellular fatty acid oxidation. Intracellular fatty acids are converted by different enzymes including DGAT2 into triacylglycerols(TAG), which are sequestered into lipid droplets. *M. tuberculosis* gains access to host derived fatty acids, e.g. via the interaction of bacteria containing phagosomes with TAG-rich lipid droplets. Intracellular accumulation of fatty acids also induces necrotic cell death (lipotoxicity) thereby promoting Mtb dissemination and release of lipid droplets from the dying host cell.

Independent studies have documented that pathogens including different mycobacterial species can trigger foam cell formation in a Toll-like receptor (TLR) mediated manner^56–58,59,60^. From a metabolic perspective, exposure of macrophages to already a single TLR ligand increases TAG storage^12,61^, enhances fatty acid uptake^12,62^, and diminishes mitochondrial fatty acid oxidation even in the presence of sufficient oxygen^12,62^. We have previously shown that synthetic lipopeptides (Pam3CSK4), lipopolysaccharide (LPS) and various mycobacterial species including Mtb induce WNT6 in a TLR - NF-κB-dependent manner^25^. We now demonstrate that WNT6 drives the accumulation of TAG-rich lipid droplets by inducing the expression of several key lipid metabolic enzymes involved in neutral lipid synthesis and storage including ACC2. This key regulatory enzyme is well known to promote neutral lipid storage by blocking fatty acid oxidation^37^ as it inhibits carnitine palmitoyltransferase 1 (CPT1)-dependent fatty acid uptake into mitochondria (summarized in Figure 6). Our results suggest that mycobacteria-induced and TLR-dependent differentiation of macrophages into a foamy phenotype is caused by WNT6-ACC2-induced metabolic changes in these cells. During chronic Mtb infection, mycobacterial TLR ligands lead to recurring activation of macrophages. Thus, it is likely that the TLR-WNT6-dependent perturbation of fatty acid metabolism can promote foam cell formation in pulmonary TB. Our prior findings suggest a physiological role of WNT6 in dampening inflammation^25^. This explains why this mediator is not only induced during Mtb infection but is also upregulated in various chronic inflammatory disease settings such as inflammatory bowel disease and allergic asthma^63,64^.

Functional evidence for the importance of complex lipids as nutrient source for Mtb originates from studies showing that Mtb growth is inhibited when neutral lipid accumulation is diminished^65,66^. Mtb has evolved different mechanisms to manipulate host lipid metabolism. It disturbs cholesterol homeostasis by activating cells with keto-mycolic acid^14,67^, or impairs degradation of complex lipids via the anti-lipolytic GPR109A GPCR receptor^65^. Recently, it was shown that Mtb infection reduces fatty acid oxidation of macrophages by inducing miRNA-33, which inhibits lipid degradation and promotes Mtb growth^66^. Our current study reveals the metabolic consequences of WNT6 expression in macrophages during pulmonary TB, revealing that WNT6 shifts lipid metabolism away from oxidation of fatty acids towards intracellular retention of TAGs. Further, our observation that *Wnt6*- and ACC2-deficiency, as well as pharmacological inhibition of ACC2 impairs Mtb replication in macrophages demonstrates that the WNT6-ACC2 axis is exploited by the pathogen to facilitate its intracellular growth. Our data suggest that WNT6 and ACC2 activity control the intracellular availability of TAG-derived fatty acids utilized by Mtb in the macrophage. This is due to the fact, that (i) addition of fatty acids rescues Mtb growth in the absence of both functional WNT6 and ACC2 and (ii) inhibition of ACC2 limits incorporation of host-derived oleic acid into Mtb specific cell membrane phospholipids. These findings are consistent with observations on Mtb’s ability to utilize oleate-induced host lipid droplets as carbon source^20^, and with data showing that proteins of Mtb, which are critically involved in fatty acid transport ^68,69^, are required for full virulence of Mtb *in vivo*^68^. It has also been shown that – when macrophages face hypoxic conditions - host-derived fatty acids are converted by Mtb into TAGs that are stored in the bacteria^70^. Fatty acids from Mtb-TAGs may not be used for bacterial replication under these conditions. Ultimately, the microenvironment of Mtb and its host cell - in particular local oxygen levels - are decisive whether Mtb can actively replicate or acquires a dormancy-like phenotype.

Lipid droplets are multifunctional organelles, which consist of proteins, enzymes and various types of neutral-lipids such as TAGs and CEs ^71,72^. Recent findings show that also cytokines drive the accumulation of lipid droplets in the context of experimental TB infections^73^. If one relates the data by Knight et al.^73^ to the results presented here, it appears that Interferon-gamma/HIF1-a signaling mediated lipid droplet accumulation is largely dependent on CE synthesis^73^. In contrast, we found that, in the absence of IFN-g, the Mtb-induced WNT6-ACC2 signaling pathway drives the accumulation of TAG-rich lipid droplets in macrophages. It is possible that during Mtb infection, bacterial and host-derived signals induce the formation of differentially composed subsets of lipid droplets, either being rich in TAG or CE. Thus, the amount of Mtb bacteria in relation to the extent of the host response may define whether TAG or CE rich lipid droplets are formed. Depending on their composition, lipid droplets could either contribute to host defense by acting as a platform for the synthesis of small lipid mediators^73^ or rather promote bacterial replication by being exploited by Mtb as a carbon source. Our current study findings suggests a unique role for WNT6-ACC2 inducible TAG-rich lipid droplets in promoting Mtb replication during infection.

In post-primary TB granulomas, necrosis is often associated with functional disintegration of the structured tissue reaction, which ultimately causes rupturing into the airways and dissemination of the mycobacteria into adjacent cells. The caseum, the liquefied content of necrotized granulomas, consists of host-derived lipids including TAGs^15^ indicating that foamy macrophages undergo cell death during granuloma progression. Cell death is a multifaceted process induced by a variety of mediators^74^. Among these an intracellular accumulation of fatty acids has been shown to exert lipotoxic effects on cells^75^. Importantly, it has been shown that fatty acid induced cellular toxicity is diminished by inhibition of ACC^46^. Consistent with this observation, we demonstrate that Mtb-infected human macrophages show significantly reduced rates of necrotic cell death when ACC2 is inhibited by pharmacological inhibitors. Moreover, we observed that ACC2 inhibition during experimental TB infection of TB susceptible mice reduces the expression of granuloma disintegrating and necrosis-inducing factors, as the formation of chemokines (CXCL1 and 5) in inhibitor treated mice are significantly reduced. Consequently, WNT6-ACC2 signaling not only induces foam cell formation during pulmonary TB, but may also contribute to subsequent necrotic cell death of foamy macrophages and granuloma disintegration, thus favoring infection of neighboring cells and the dissemination of bacteria within and from the host. In contrast to apoptosis^74^, necrotic cells have been suggested to serve as niches for Mtb replication^49,50^ and represent a way for Mtb to disseminate extracellulary^48^.

While more than 10 drugs are currently available for TB treatment, treatment success of MDR and XDR Mtb strains is as low as 50% on a global level. This stimulated intensive research to develop new anti TB drugs, but also to explore alternative treatment concepts including host-directed therapies (HDT), which bear the promise of enhancing the efficacy of classical TB drugs and prevent resistance development^2^. Targeting the formation of TAG-laden “foamy” macrophages in the host may represent a promising HDT approach, as our findings suggest that these cells are exploited by intracellullar Mtb to access lipids as predominant carbon source. A deregulated cellular lipid metabolism, as observed in the development of diseases such as hepatic steatosis^39^ and non-small cell lung cancer^52^, can successfully be treated *in vivo* by the use of ACC2 inhibitors. Our findings demonstrate that pharmacological interference with WNT6-ACC2 signaling indirectly targets Mtb (i) by depriving Mtb from TAG-derived nutrients within the intracellular niche, and (ii) by limiting Mtb-induced necrotic cell death and mycobacterial dissemination. Our current *in vivo* data indicate that targeting ACC2 could complement standard TB therapy as ACC2 inhibition in antibiotic-treated mice reduced the abundance of nutritive lipids and reduced the release of pro-necrotic mediators. Additional animal studies are necessary to further improve the concomitant therapy with ACC2 inhibitors when INH is administered. This includes the optimization of dosage, the start and the duration of therapy. In a long term pesrpective targeting of the host metabolic enzyme ACC2 may represent a promising approach for a host-directed adjunct therapy of antibiotic-based TB treatment.

## Supplementary figure legends

**Figure S1:**
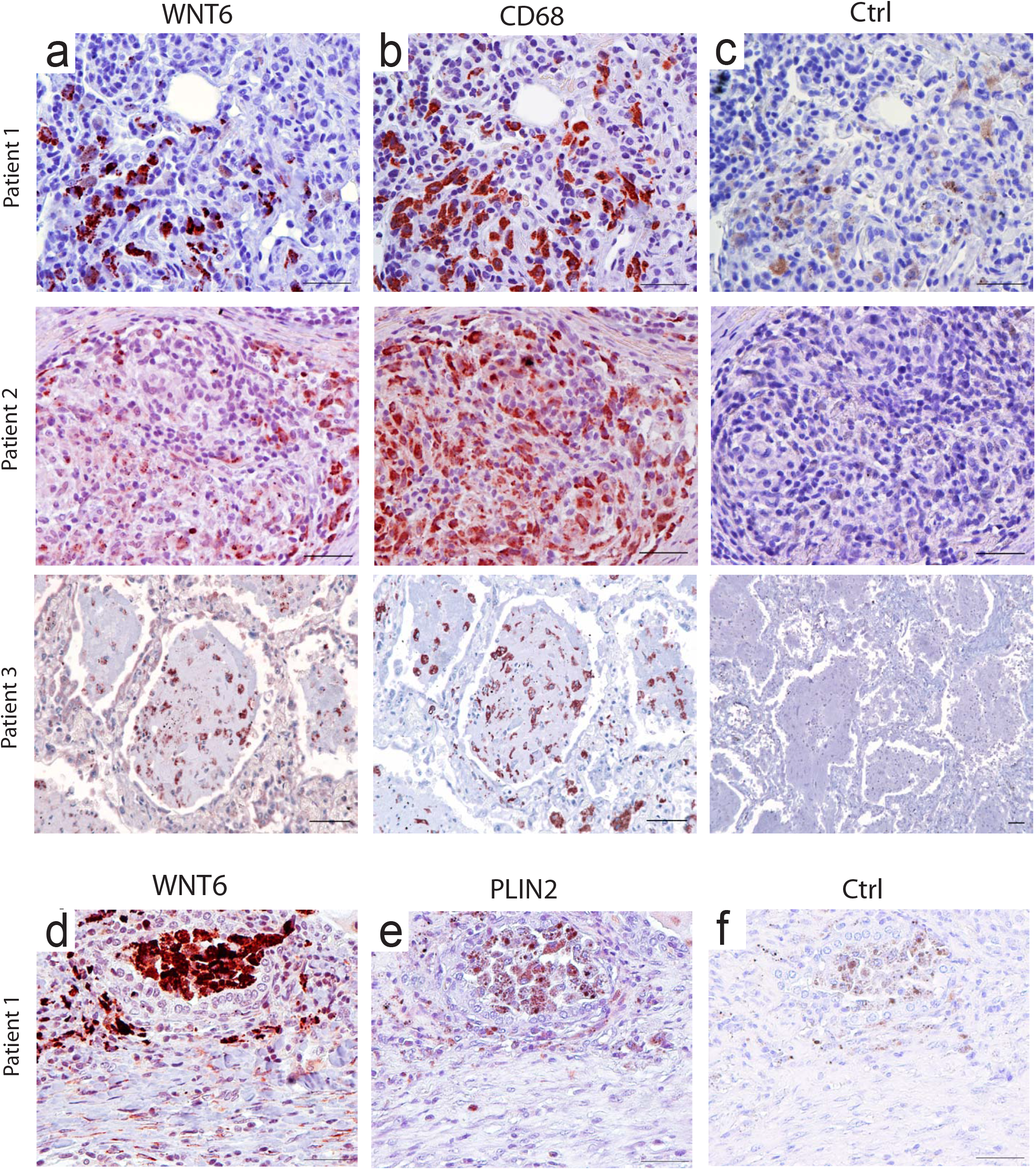
WNT6 expression in lungs of TB patients. Immunohistochemical analyses of formalin-fixed and paraffin-embedded lung tissue from three tuberculosis patients. Sections (1-2 μm) were incubated with antibodies specific for WNT6 **(a, d)**, the macrophage/monocyte marker CD68 **(b),** the lipid droplet scaffolding protein Perilipin 2 (ADFP) **(e)** or without primary antibodies as a control **(c,f)**. Antigens were visualized with a horseradish-peroxidase (HRP)-based detection system using AEC as chromogen. Scale bar: 50 μm and 100 μm (c (ctrl of patient 3)).

**Figure S2:**
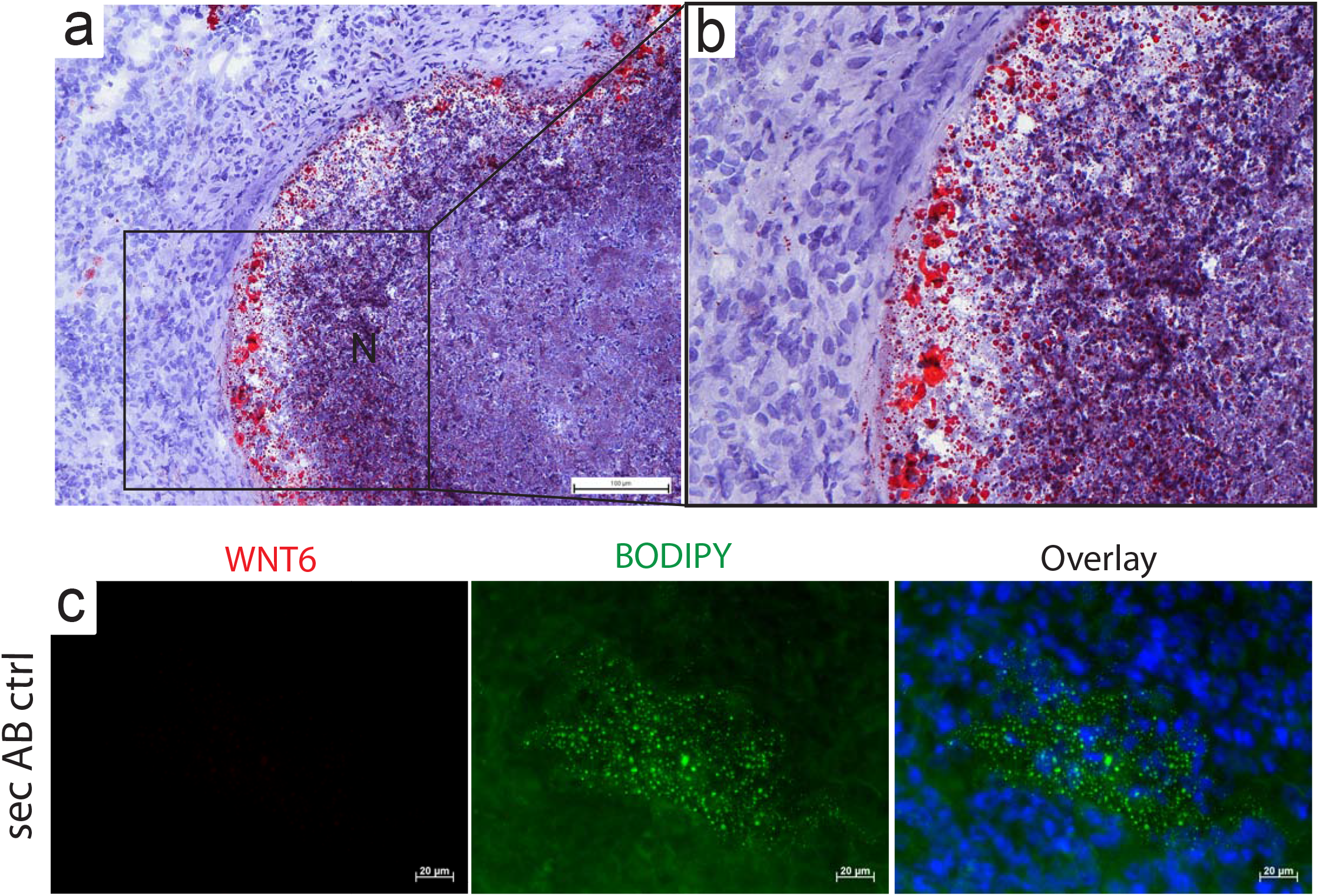
WNT6 and neutral lipids in Mtb-infected *IL-13* overexpressing mice. Frozen lung tissue sections (5μm) derived from Mtb-infected *IL-13* overexpressing mice were stained for neutral lipids with Oil-Red O (**a, b;** d104 p.i.) or BODIPY 493/503 (**c**; d63 p.i.,10 μg/ml, green). The section in (c) was stained with BODIPY in the absence of primary antibody (sec. AB ctrl.). Nuclei were stained with DAPI (blue); N, Necrosis.

**Figure S3:**
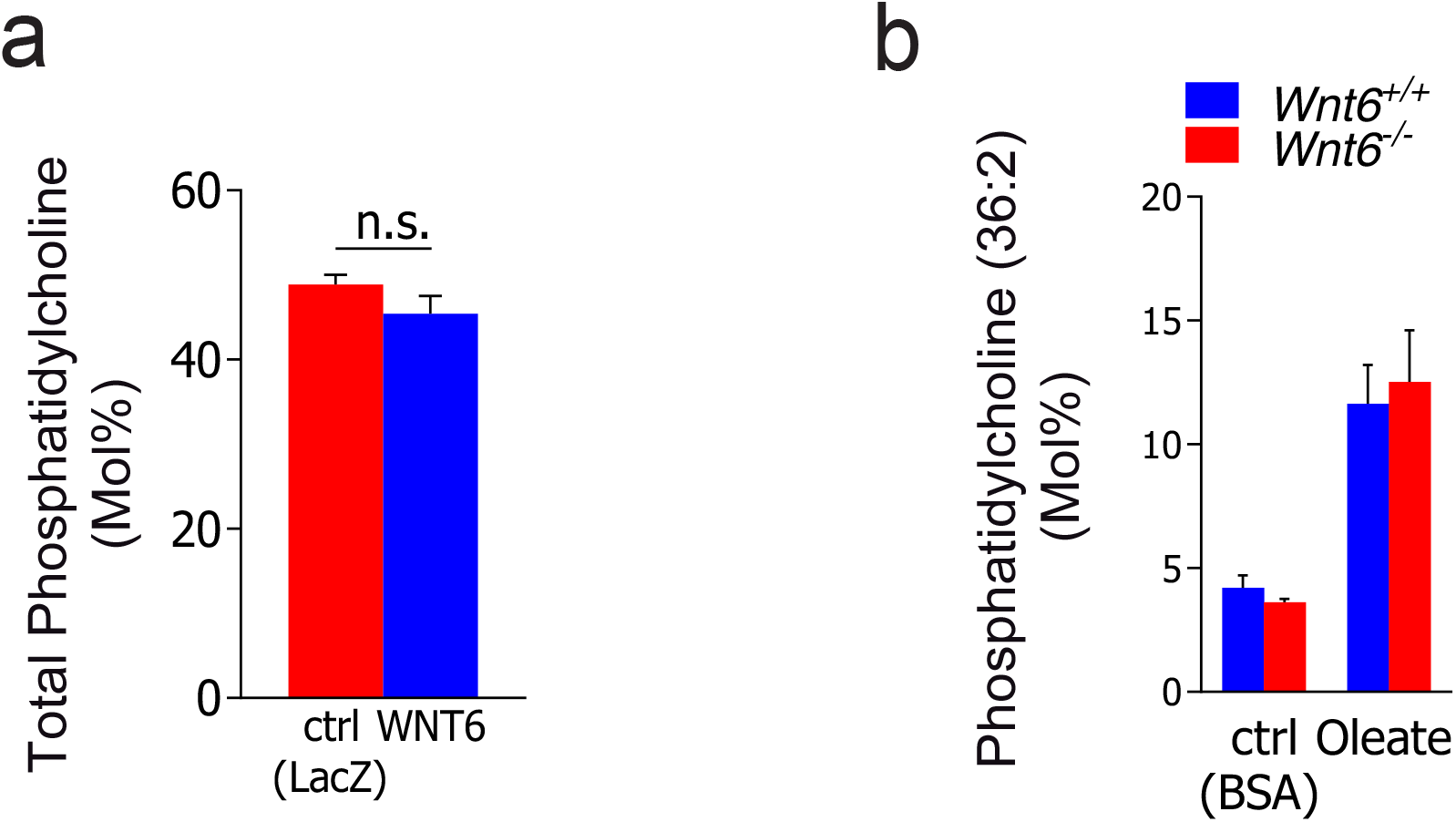
WNT6 does not affect synthesis of phosphatidylcholines. **(a)** Quantification of phosphatidylcholine species in WNT6-overexpressing or control (LacZ) NIH3T3 cells by mass spectrometry. The sum of all measured phosphatidylcholine species (expressed in Mol %) from the same set of experiments as depicted in Figure 2c is shown; n=3. **(b)** Phosphatidylcholine 36:2 levels in *Wnt6*^+/+^ or *Wnt6*^−/−^ BMDMs incubated for 24 hours in the absence (BSA, ctrl) or presence of fatty acids (oleate-BSA, 200 μM) as determined by mass spectrometry. Data are from the same set of experiments as depicted in Figure 2e; n=2. All data are depicted as mean +/− SEM.

**Figure S4:**
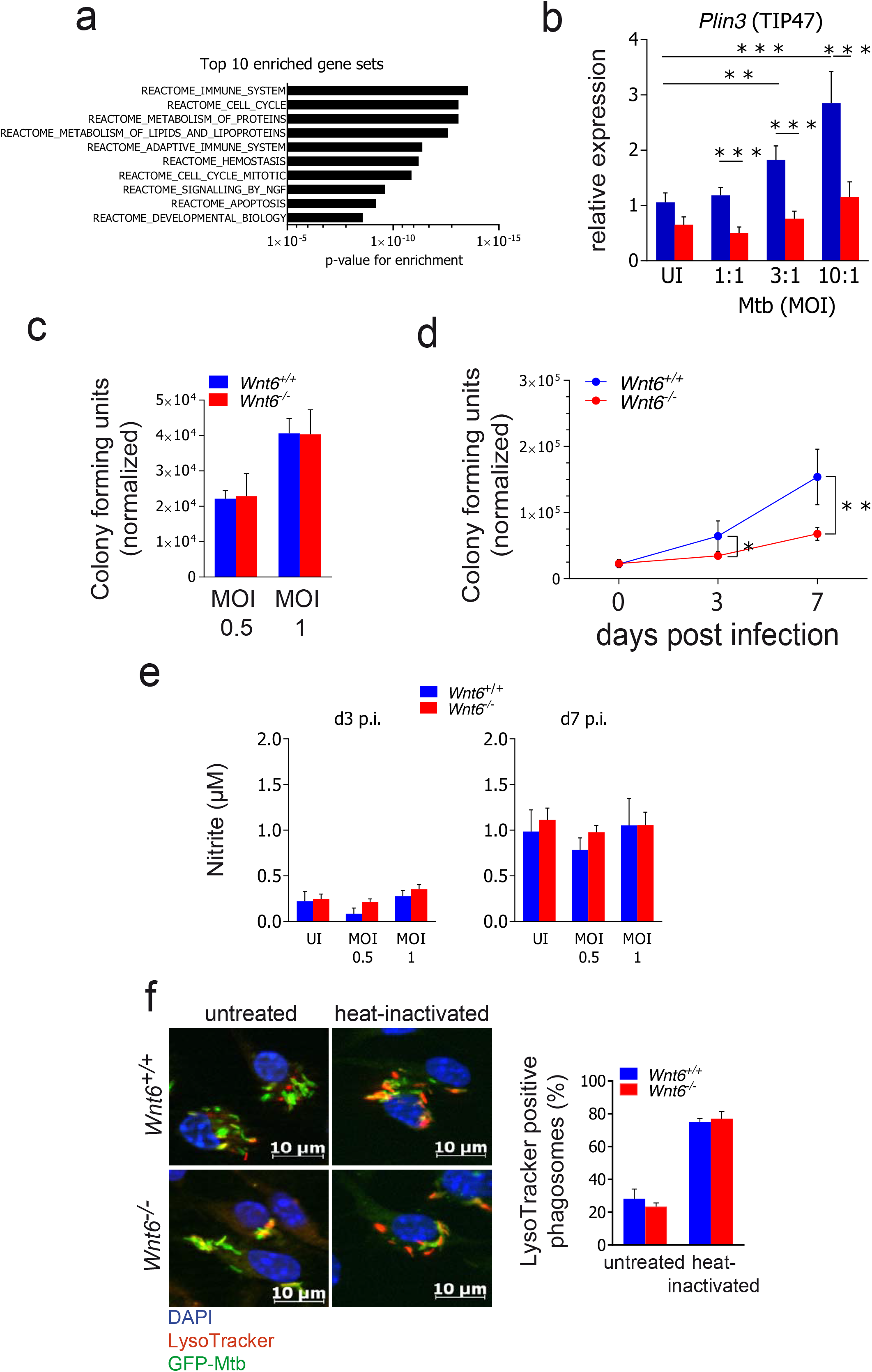
Effect of *Wnt6*-deficiency on Mtb-induced gene expression, Mtb uptake and growth, nitrite formation and acidification of Mtb-containing compartments. **(a)** Top 10 enriched gene sets derived from a gene set enrichment analysis of all differentially regulated genes between Mtb-infected *Wnt6*^+/+^ and *Wnt6*^−/−^ BMDMs. Cells were infected for 24 h with Mtb H37Rv (MOI 3:1), total RNA was extracted, and subjected to microarray-based gene expression analyses. Further analysis was conducted as described in *Material and Methods*;n=3. **(b)** qRT-PCR based gene expression analysis of *Wnt6*^+/+^ and *Wnt6*^−/−^ BMDMs infected for 24h with various doses (MOIs) of Mtb H37Rv; n=3. **(c,d)** CFU analysis of *Wnt6*^+/+^ and *Wnt6*^−/−^ BMDMs after infection with Mtb H37Rv. Cells were infected, washed (4 hours p.i.) and incubated for the time indicated. Bacterial growth was related to the number of macrophages (normalized CFU) at the individual timepoint (given as CFU per 100.000 cells); n=3. **(e)** Quantification of nitrite (NO ^−^) in culture supernatants of *Wnt6*^+/+^ and *Wnt6*^−/−^ BMDMs after infection with Mtb H37Rv (as described in (d)) for 3 (left panel) or 7 days (right panel) by Griess reaction; n=2. **(f)** Visualization (left panel) and quantification (right panel) of acidified Mtb-containing compartments. *Wnt6*^+/+^ and *Wnt6*^−/−^ BMDMs were infected with heat-inactivated (85°C, 5 minutes) or viable GFP-expressing Mtb H37Rv (green) for 4 hours, were simultaneously (2 h) treated with LysoTracker dye (400 nM; red), washed, fixed, stained with DAPI (1 μg/ml; blue) and visualized by fluorescence microscopy. Evaluation of LysoTracker positive phagosomes was conducted in a blinded fashion of over 200 compartments per condition in a total of 3 independent experiments. Statistical analyses were carried out using One-Way ANOVA with a suitable post-hoc test for multiple comparison (d); *p≤0.05, **p≤0.01, ***p≤0.001. All data are depicted as mean +/− SEM.

**Figure S5:**
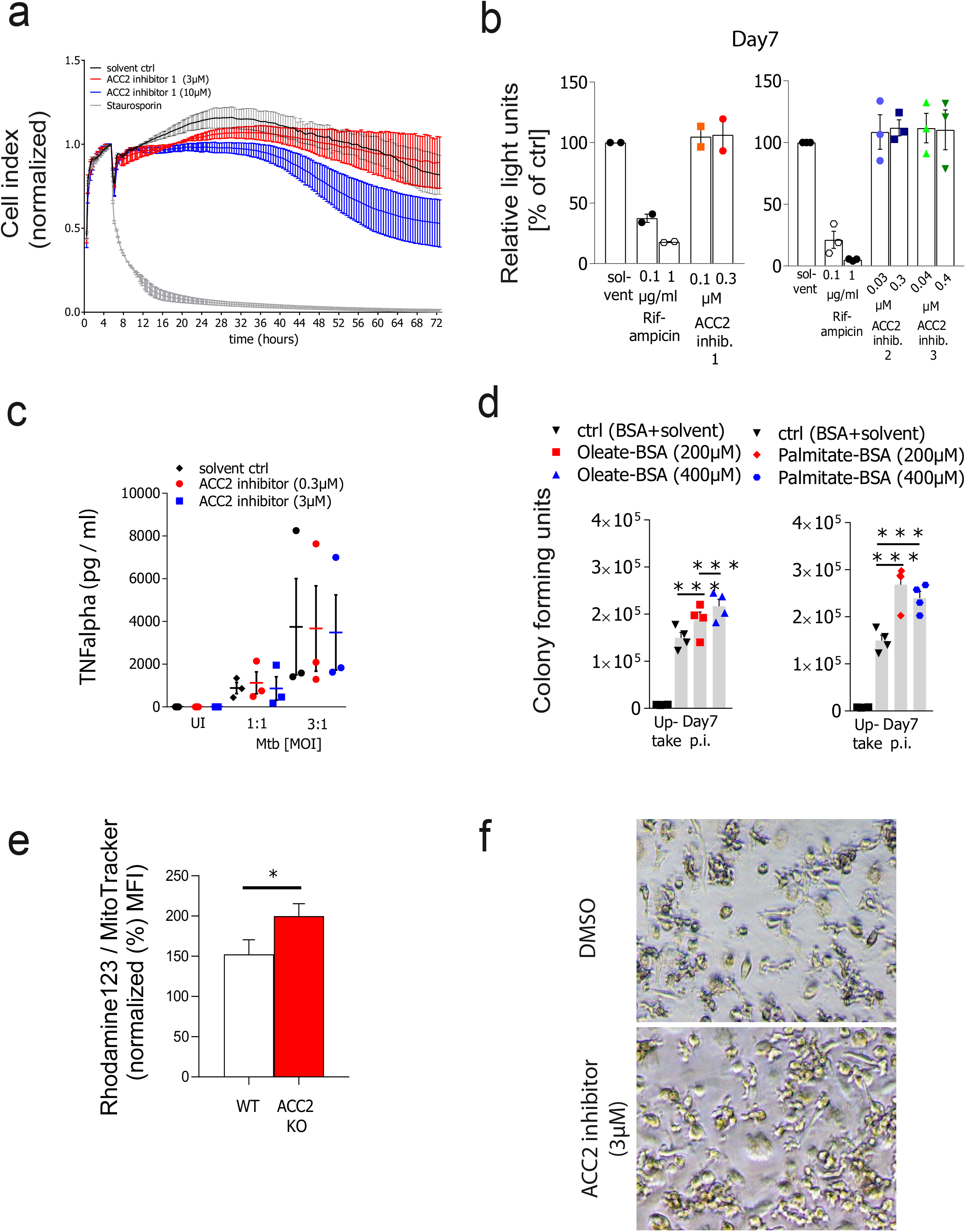
Pharmacological inhibition of ACC2 does not decrease replication of Mtb in liquid culture, neither does it decrease viability or affect TNFα release of Mtb-infected primary human macrophages. **(a)** Analysis of human macrophage viability in the presence of ACC2 inhibitor 1 as determined by real-time impedance measurements (expressed as cell index). hMDMs were incubated in the presence of solvent (DMSO, ctrl), ACC2 inhibitor 1 or Staurosporine (1 μg/ml) for the indicated time on a xCELLigence System; Depicted is representative data from 2 independent experiments with similar results. **(b)** Mtb growth in the absence (solvent) and presence of various ACC2 inhibitors or the TB drug rifampicin as determined by measuring fluorescence of GFP-expressing Mtb in liquid culture. Bacteria were cultured in 7H9 medium supplemented with 10% OADC and growth was measured as relative light units at 528 nm after excitation at 485 nm in a fluorescence microplate reader at the indicated time point; n=2 (left panel), n=3 (right panel). **(c)** TNFα release of hMDMs infected for 24 hours with Mtb H37Rv and simultaneously incubated with solvent (DMSO, ctrl) or the indicated concentrations of ACC2 inhibitor 1. Mean +/− SEM from 3 independent experiments/donors is shown. **(d)** Effect of addition of fatty acids on Mtb CFU in primary human macrophages. After infection with Mtb (MOI 0.5:1), cells were washed and incubated in the absence (BSA) or presence of different concentrations of oleate- and palmitate-BSA. Data are derived from the same set of experiments shown in Figure 4f; n=4. **(e)** Flow cytometry-based quantification of Rhodamine 123 signals (relative to MitoTracker Deep Red signals (both aMFI) x100) in Mtb-infected wild-type (WT) and ACC2 KO human macrophage-like cells (BLaER1 macrophages) (MOI 0,1:1) normalized to uninfected WT cells at day 3 p.i‥**(f)** Enhanced viability of hMDMs during infection with Mtb H37Rv when treated with ACC2 inhibitor (lower panel) in comparison to solvent control (DMSO, upper panel). Depicted is a representative observation of 2 independent experiments with similar results. Statistical analyses were carried out using One-Way ANOVA with a suitable post-hoc test for multiple comparison (d); * p≤0.05, ***p≤0.001. All data are depicted as mean +/− SEM.

**Figure S6:**
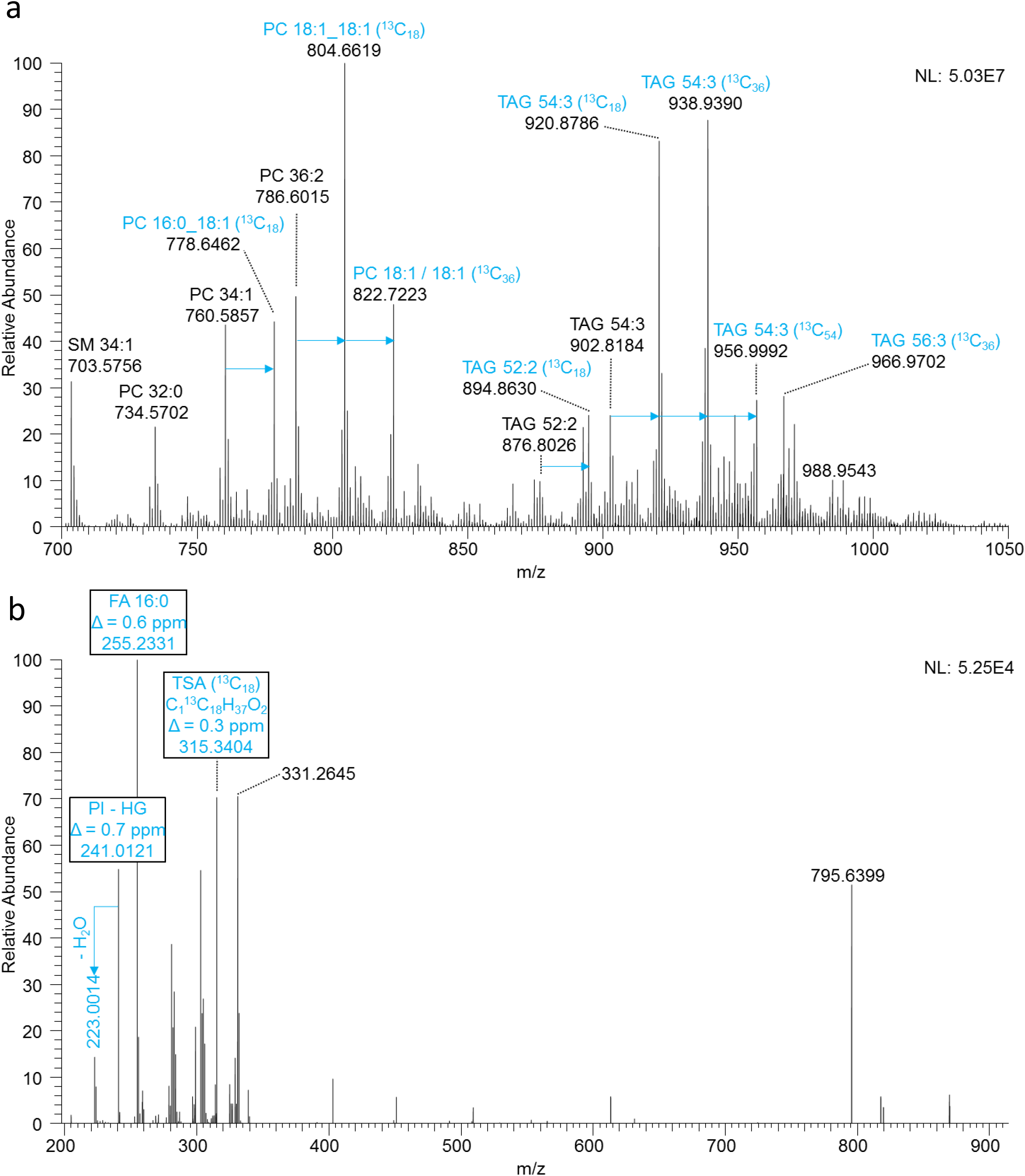
Incorporation of ^13^C-oleic acid into lipids of macrophages and metabolization to tuberculostearic acid (TSA) to form phosphatidylinositol (PI) 16:0_19:0 (TSA) of *Mtb*. Mass-spectometric analysis of ^13^C-oleate pulsed and Mtb-infected hMDMs showing **(a)** incorporation of oleic acid into major abundant lipids of macrophages. The mass shift of ^13^C labelled lipid species is shown in blue as determined in positive ion mode MS^1^. Data are from the same set of experiment as shown in Figure 4g and h. **(b)** Tandem mass spectrometric analysis of ^13^C labelled PI 16:0_19:0 (TSA) with the precursor m/z 869.6 in the negative ion mode. Specific fragment ions for identification of the lipid are shown in blue (PI - HG: fragments ion of the phosphatidylinositol head group, TSA (^13^C18): isotopically labelled tuberculostearic acid with incorporation of one ^12^C methyl group as described (Heyckendorf et al.; Biorxiv, 2020). Data are from the same set of experiment as shown in Figure 4g and h and originate from infected macrophages of donor 2 after 7 days of infection (Supplementary Table I).

**Figure S7:**
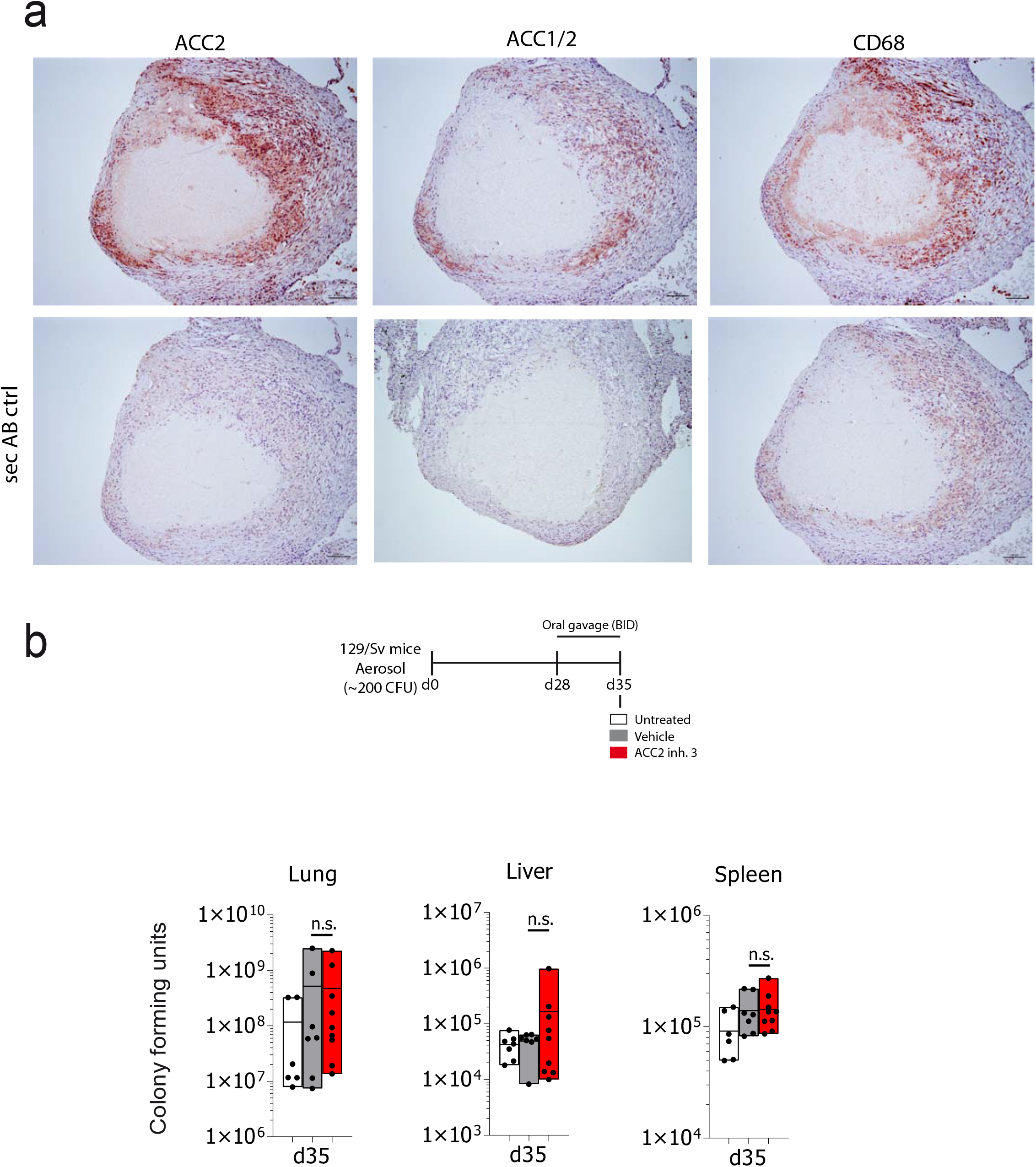
ACC2 is expressed in human lung tissue of TB patients and its inhibition by a low-dose and short-term treatment with a pharmacological inhibitor does not affect bacterial replication in Mtb-infected mice. **(a)** Immunohistochemical analyses of formalin-fixed and paraffin-embedded lung tissue derived from a tuberculosis patient (Patient 1). The upper panel shows consecutive sections (1 μm) incubated with primary antibodies directed against ACC2 (left panel), ACC1/2 (middle panel) and the macrophage/monocyte marker CD68 (right panel). The lower panels show the respective consecutive section, which was incubated without primary antibody (sec. AB ctrl). Antigens were visualized with a Horseradish peroxidase (HRP)-based detection system using AEC as chromogen (red). **(b)** Effect of a low-dose and short-term ACC2 inhibitor treatment on mycobacterial loads in Mtb-infected mice. After 28 days of infection with a low dose of Mtb H37Rv (~200CFU), 129/Sv mice were either left untreated (white bars), were treated with vehicle solution (grey bars) or with ACC2 inhibitor (ND-646, 25 mg/kg BW) for a period of 7 days. At day 35 p.i., Mtb bacterial burden was determined in lung, liver and spleen (n=6-10 animals per group). Statistical analyses were carried out using an one-tailed, unpaired Student’s t-test; n.s.= not significant. Data are depicted as Min-Max bar with line at mean.

## Acknowledgements & Funding

The authors are very grateful for the funding within the DFG priority program (SPP1580) (NR: Re1228 5-1, Re1228 5-2), the Cluster of Excellence 306 (“Inflammation at interfaces”), and the Deutsches Zentrum für Infektionsforschung (DZIF) within the “Thematic translational unit tuberculosis” (TTU TB; CH: TTU 02.705; NR: TTU 02.806; 02.810; DS: TTU 02.704-1, 02.811). Moreover, we would like to gratefully acknowledge Carolin Golin, Lisa Niwinski and Johanna Volz for expert technical assistance.

## Competing interests

Drs. N. Reiling and J. Brandenburg (Research Center Borstel, Leibniz Lung Center, 23845 Borstel, Germany) have filed a patent application entitled “ACC inhibitors as means and methods for treating mycobacterial diseases”(WO2018007430A1, patent pending).

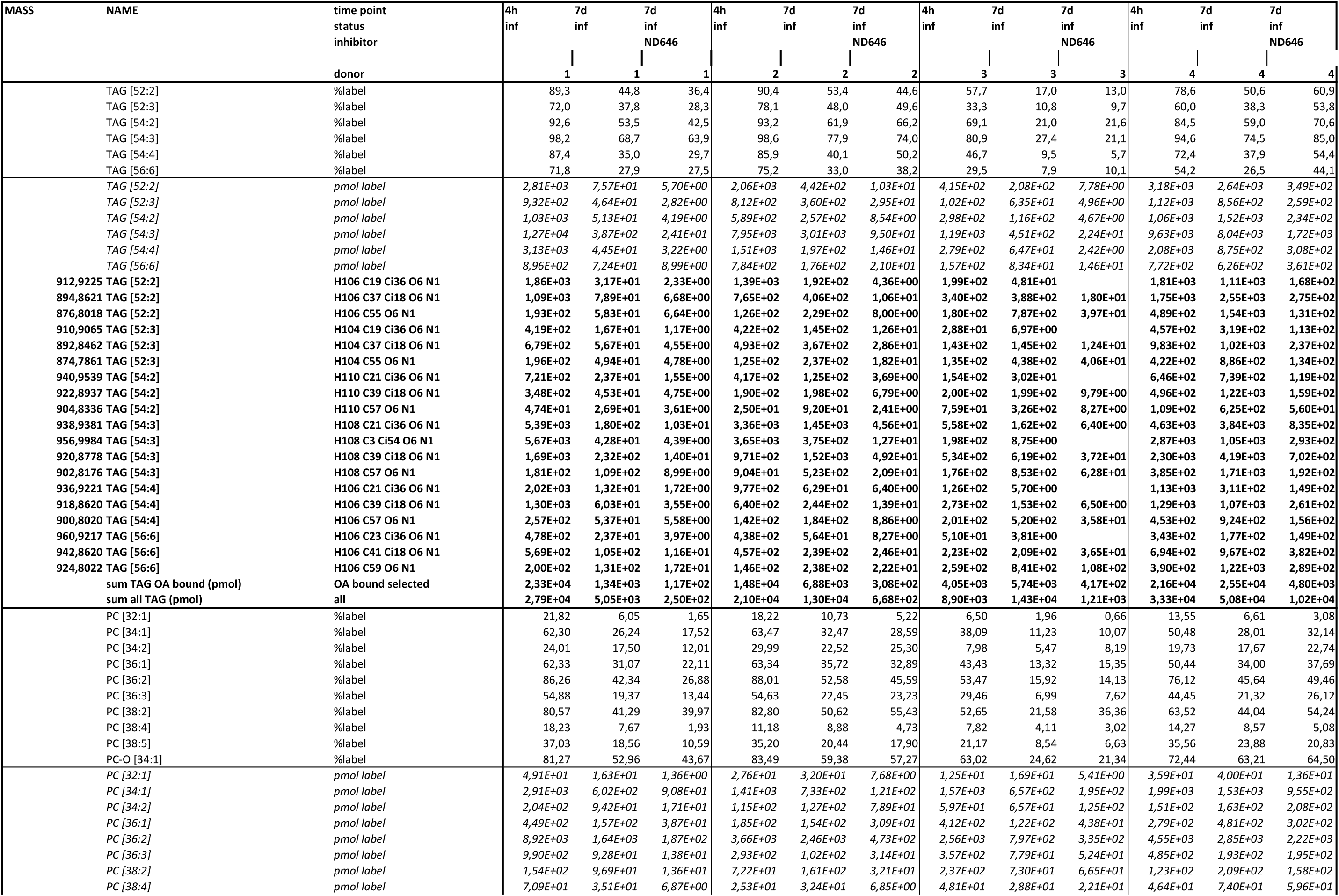
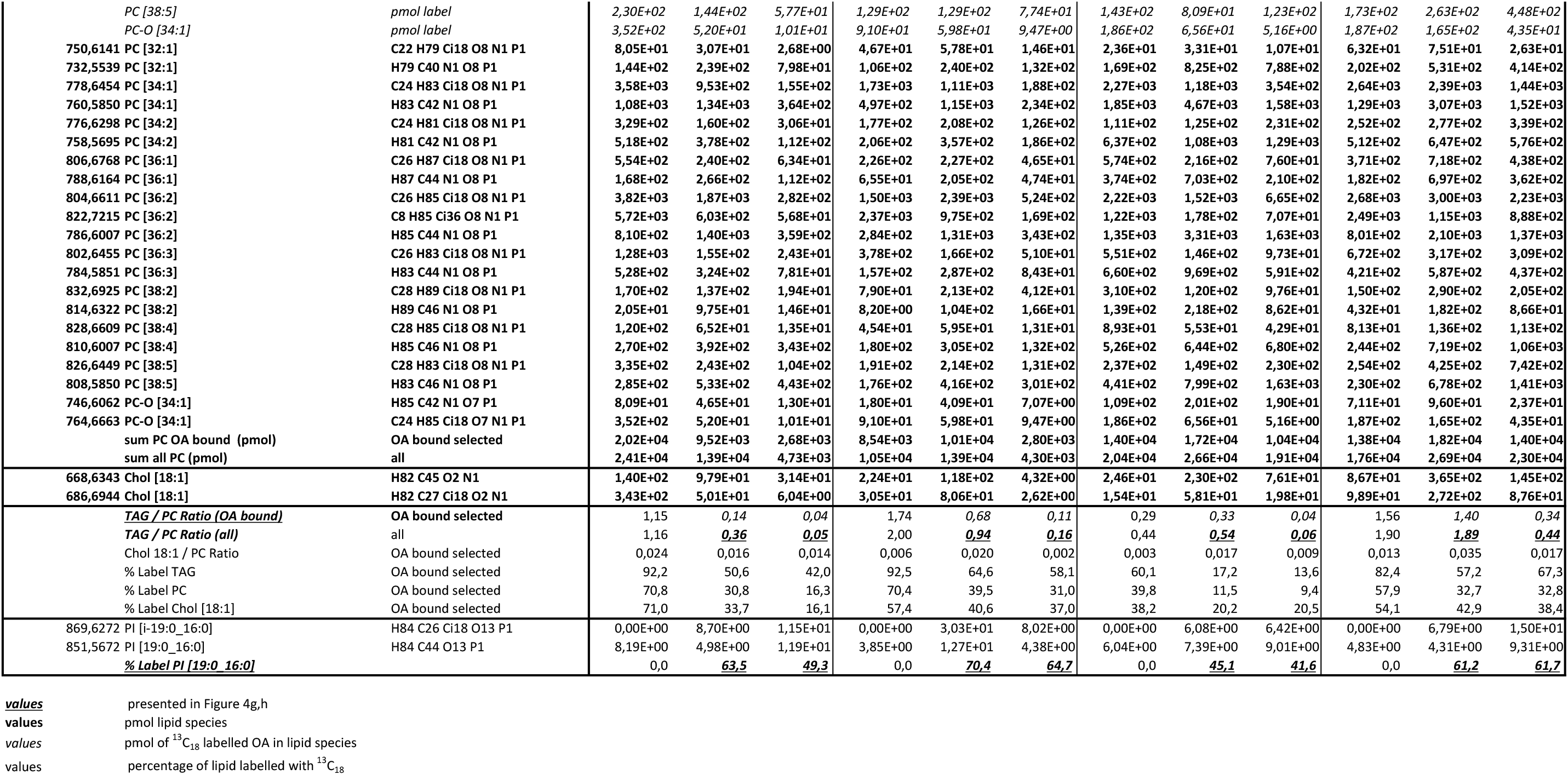
Summary for Tracer Analysis of ^13^C_18_ labelled Oleic Acid during *Mtb* Infection of Human Monocyte‐Derived Macrophages (hMDM)

## Materials and Methods

### Mice and macrophages

129/Sv mice were purchased from Janvier (Le Genest-Saint-Isle, France). NMRI *Wnt6*^+/+^ or *Wnt6*^−/−^ and *IL-13*-overexpressing mice were raised and maintained under specific pathogen-free conditions. The Wnt6 null allele was generated as described previously^77^. NMRI *Wnt6*^−/−^ mice were generated by heterozygous mating at the Research Center Borstel. *IL-13* overexpressing mice were generated as described elsewhere^78^ and kindly provided by Andrew McKenzie (Cambridge, UK).

To generate bone-marrow derived macrophages (BMDM), mice were sacrificed, and bone-marrow cells were flushed out from femora and tibiae with ice-cold DMEM as described previously^79^. To yield high purity and remove contaminating fibroblasts^80^, bone-marrow cells were first cultivated in Nunclon Delta cell culture dishes (Thermo Fisher, Waltham, USA) for 24 hours. Only non-adherent cells were collected, and incubated for 7 days in cell culture dishes (Sarstedt, Nümbrecht, Germany) in DMEM containing 10 mM HEPES, 1 mM sodium pyruvate, 4 mM glutamine (Biochrome, Berlin, Germany), 10% of heat-inactivated fetal calf serum (FCS; Pan-Biotek, Aidenbach, Germany), supplemented with 50 ng/ml macrophage colony stimulating factor (M-CSF; Bio-Techne, Minneapolis, USA)^81^. To obtain quiescent tissue macrophages^82^, peritoneal exudate cells (PEC) were isolated from the resting peritoneal cavity of mice as described previously^79^. To generate human monocyte-derived macrophages (hMDM), peripheral blood monocytes (purity consistently >92%) were obtained by counterflow centrifugation from peripheral blood mononuclear cells (PBMCs) of healthy blood donors. Subsequently, isolated cells were incubated for 7 days in Teflon bags (VueLife 72C; Cellgenix, Freiburg, Germany) in VLE RPMI 1640 (Biochrome) containing 4% human AB serum, 4 mM glutamine, 1% penicillin/streptomycin (Merck, Darmstadt, Germany) and 10 ng/ml recombinant human M-CSF as described previously^83^. All cells were incubated in cell culture medium with the omission of M-CSF before proceeding further.

### *M. tuberculosis* strains and *in vitro* growth assays

*M. tuberculosis* strain H37Rv (ATCC 27294; American Type Culture Collection, Manassas, VA), GFP-expressing *M. tuberculosis* (H37Rv::pMN::437^84^ or H37Rv::psVM4^85^), and mCherry-expressing *M. tuberculosis*^86^ were harvested at mid-log phase (OD_600nm_ ~0.3) and stored as frozen aliquots at −80°C as described previously^83^.

For *M. tuberculosis* growth analysis in liquid culture, frozen aliquots were thawed, centrifuged (2300×*g*, 10 minutes) and bacteria in 7H9 medium supplemented with 10% Oleic Albumin Dextrose Catalase (OADC) (Sigma, St.Louis, USA) thoroughly homogenized by use of a syringe and a 26-gauge syringe needle. Subsequently, 2 × 10^6^ bacteria were cultured in a total volume of 100 μL in a black 96-well plate with a clear bottom (Corning, New York, USA) and were sealed with an air-permeable membrane (Porvair Sciences, Wrexham, UK). Growth was measured as relative light units at 528 nm after excitation at 485 nm in a fluorescence microplate reader (Synergy 2, BioTek Instruments, Vermont, USA) at the indicated time points.

### Infection of macrophages and mice

For *in vitro* infection experiments, Mtb bacteria from frozen aliquots were homogenized as described above and resuspended in cell culture medium. Cells were infected with the indicated dose of bacteria (multiplicity of infection (MOI)) and, if not indicated otherwise, incubated for 4 hours (37°C, 5% CO_2_), followed by extensive washing with Hanks Buffered Salt Solution (HBSS, Sigma) in order to remove extracellular bacteria. Subsequently, cells were treated with solvent/carrier control, the indicated inhibitor or fatty acids for up to 7 days (37°C, 5% CO_2_).

For quantification of viable colony forming units (CFU) in macrophages, cells were lysed by incubation with 2% Saponin in HBSS and lysates were serially diluted in 0.05% Tween-80 / dH_2_0 and plated on 7H10 agar plates containing 10% heat-inactivated bovine serum (Merck, Darmstadt, Germany). Plates were incubated for 3 to 4 weeks at 37°C. Before lysing cells, images were taken at defined positions of each well by use of a bright field microscope (DM LB, Leica Biosystems, Wetzlar, Germany) and a digital camera (Sight DS-L11, Nikon, Tokio, Japan). The number of cells within a well was enumerated by analyzing images with a counting tool (Adobe Photoshop CS5 software, Version 12.04 and earlier). The Mtb/macrophage ratios were calculated at the individual time point based on the obtained CFU data and the enumerated number of cells per well.

C57BL/6, 129/Sv and *IL-13* overexpressing mice were infected via the aerosol route with *M. tuberculosis* H37Rv (see above) as described previously^26,87^. During infection experiments, mice were kept under barrier conditions in the biosafety level 3 facility at the Research Center Borstel in individually ventilated cages. For analysis of lung bacterial loads, lungs from sacrificed animals were removed aseptically, weighed and homogenized in PBS containing a proteinase inhibitor cocktail (Roche Diagnostics, Mannheim, Germany) using the FastPrepTM System (MP Biomedicals, Solon, USA). Tenfold serial dilutions of organ homogenates were plated onto Middlebrook 7H10 agar plates containing 10 % heat-inactivated FBS. After an incubation at 37°C for 21 days, colonies on plates were enumerated. All animals were weighed regularly before and after infection using a laboratory balance, and as a means for evaluating disease progression the body weight change was calculated.

### Stimuli and inhibitors

For *in vitro* infection experiments, dimethylsulfoxid (DMSO for cell culture; Sigma) was used to solubilize N-(1-(2′-(4-Isopropoxyphenoxy)-2,5’-bithiazol-5-yl)ethyl)acetamide (“ACC2 inhibitor 1”; ab142090; purchased from Abcam, Cambridge, UK), 5-[1’-(1-cyclopropyl-4-methoxy-3-methylindole-6-carbonyl)-4-oxospiro[3H-chromene-2,4’-piperidine]-6-yl]pyridine-3-carboxylic acid (“ACC2 inhibitor 2” known as MK-4074^39^; purchased from MedChemExpress, Sollentuna, Sweden) and 1,4-dihydro-1-[(2R)-2-(2-methoxyphenyl)-2-[(tetrahydro-2H-pyran-4-yl)oxy]ethyl]-a,a,5-trimethyl-6-(2-oxazolyl)-2,4-dioxothieno[2,3-d]pyrimidine-3(2H)-acetamide (“ACC2 inhibitor 3” known as ND-646^52^; MedChemExpress, Sollentuna, Sweden). DMSO served as a solvent control (0.1% in cell culture medium).

^12^C-Oleic acid (pure, pharma grade; Applichem, Munich, Germany), and ^12^C-Palmitic acid (Sigma) were conjugated to the carrier protein Bovine Serum Albumin (BSA; Applichem or Sigma (low-endotoxin, fatty-acid free)) according to the protocol of Listenberger et al^88^. Briefly, a solution of 20 mM fatty acid in 0.01 M NaOH was incubated at 70°C for 30 minutes, followed by dropwise addition of 1 M NaOH facilitating the solubilisation of the fatty acid. Solubilised fatty acids were complexed to BSA in PBS at a 8:1 fatty acid to BSA molar ratio. The complexed fatty acids or BSA alone were added to serum-containing cell culture medium to achieve different fatty acid concentrations or a suitable control. Inhibitors and fatty acids were added to the cells after removing extracellular bacteria by washing in order to avoid interference with bacterial uptake.

### ACC2 inhibitor treatment of mice

To study the effect of ACC2 inhibition on Mtb infection *in vivo*, ACC inhibitor 3 (ND-646)^52^ was administered by oral gavage twice a day (BID)) at a concentration of 25 mg/kg bodyweight (BW). The corresponding volume of a vehicle solution (0.9% NaCl / 1% [v/v] Tween-80 / 30% [w/v] Captisol (CyDex Pharmaceuticals, San Diego, USA)) with the omission of ND-646 served as treatment control. Moreover, mice were treated either with isoniazid alone (10 mg/kg BW, Sigma) or as a combination of isoniazid with ACC2 inhibitor. Treatment was started at day 28 p.i. and conducted for a period of 7 days (Vehicle vs. ACC2 inhibitor) or for 14 days with isoniazid and isoniazid plus ACC2 inhibitor.

### NIH3T3 cells and generation of WNT6 conditioned medium

*Wnt6*-transfected NIH3T3 cells were a kind gift of Prof. S. Vainio (University of Oulu, Oulu, Finland). In order to yield highly pure WNT6 expressing clones, single cells were placed in 96-well plates using a FACSAria IIu cell sorter (Becton Dickinson (BD), Franklin Lake, USA) with an automated cell deposition unit (ACDU). The resulting clones were screened for WNT6 expression and selected accordingly. Control-transfected (LacZ) NIH3T3 cells were a kind gift of Prof. R. Kemler (Max-Planck-Institute for Immunobiology and Epigenetics, Freiburg, Germany). To generate conditioned medium (CM), culture supernatants of NIH3T3 cells grown for 3 days were collected, filtered through a 0.2-μm filter and stored at −80°C until further usage. CM derived from cells overexpressing and secreting WNT6 (referred to as WNT6 CM) or from a similar number of control (LacZ) cells (referred to as Control CM) were used for stimulation experiments with macrophages.

### BLaER1 cells and generation of functional protein knockouts using CRISPR/Cas9

B cell leukemia C/EBPαER clone 1 (BLaER1) cells^40^, a kind gift from Thomas Graf (Center for Genomic Regulation, Barcelona, Spain), were cultivated at a cell density between 1.5×10^5^ and 1.5×10^6^ cells / mL in VLE RPMI containing 10% of heat-inactivated fetal calf serum, 4 mM glutamine and 1% penicillin/streptomycin. In order to generate functional protein knockouts of ACC1 and ACC2, CRISPR/Cas9-mediated genome editing was used as described recently^41^. In detail, the Benchling online software (www.benchling.com, San Francisco, USA) was used to design gRNA sequences with a low off target score targeting the protein-coding regions of *ACACA* or *ACACB* gene, respectively (5’ - TTTGGGGATCTCTAGCCTAC-3’ and 5’-TAGGGAGTTTCTCCGCCGAC-3’). Oligodeoxyribonucleotides (purchased from Eurofins Genomics, Ebersberg, Germany) encoding the gRNA sequences were cloned into pU6-(BbsI)-CBh-Cas9-T2A-BFP^89^ (a kind gift from Ralf Kuehn (Max-Delbrück-Center for Molecular Medicine, Berlin, Germany) plasmid [Addgene, #64323]) using the BbSI restriction side followed by propagation of the plasmid in *E. coli* DH5α (New England Biolabs, Frankfurt, Germany). Subsequently, 1 × 10^6^ BLaER1 cells were transfected with 2 μg plasmid DNA using the Human B Cell Nucleofector Kit and Nucleofector I device (program U-15; both Lonza, Basel, Schweiz). On day 2 post transfection, BFP^+^ cells were single-cell-sorted into 96-well plates using a FACSAria IIu (BD Biosciences). After 3 weeks, DNA was isolated from the clones using the QuickExtract DNA Extraction Solution (Lucigen, Middleton, USA). Upon amplification and sequencing (Eurofins Genomics, Ebersberg, Germany) of side-specific gene stretches, the occurrence of InDels was determined using the Tracking of Indels by DEcomposition (TIDE) online software^90^. For further analysis only clones with frameshift mutations on both alleles (identified InDels for ACC1 and ACC2 KO cells were −7/+1 and −1/+2, respectively) were used, since homozygous frameshift InDels cause alterations in the protein-coding region leading to mRNA decay or generation of a nonfunctional protein^91^.

Transdifferentiation of wildtype (WT), ACC1 and ACC2 KO BLaER1 cells into macrophages was induced by cultivating cells in presence of 10 ng/ml recombinant human M-CSF, 10 ng/ml IL-3 (Peprotech, Hamburg, Germany) and 100nM β-estradiol (Sigma) for 7 days. After seeding BLaER1 macrophages onto coated (natural mussel adhesive protein, Abcam, UK) culture plates (Nunc) cells were incubated in cell culture medium in the absence of IL-3 and β-estradiol overnight before proceeding further.

### Real-time quantitative PCR

Cells of human or murine origin (0.2-1×10^6^) were lysed in Trizol (peqGOLD TriFast™; VWR International, Radnor, USA) and total RNA was extracted by use of the DirectZol® RNA MiniPrep (Zymo Research, Irvine, CA, USA) according to the manufacturer’s instructions. For reverse transcription of isolated RNA, the Maxima First Strand cDNA Synthesis Kit for real-time quantitative PCR (RT-qPCR; Thermo Fisher) was used. Gene-specific primer pairs and TaqMan probes (Universal Probe Library (UPL), Roche Applied Science, Mannheim, Germany) were designed with the UPL assay design center (ProbeFinder Version 2.45 and earlier versions; sequences and probes are given in Table I. RT-qPCR was performed using the LightCycler 480 Probe Master Kit and the LightCycler 480 II system (Roche Applied Science) as described previously^92^. Crossing point values of target and reference gene (hypoxanthine-guanine phosphoribosyltransferase, HPRT) were determined by the second derivative maximum method. Relative gene expression was calculated with the E-Method^93^ considering the individual efficiency of each PCR setup determined by a standard curve or, if this was not possible, by the 2^−ΔΔ*CT*^ method^94^.

**Table I.**
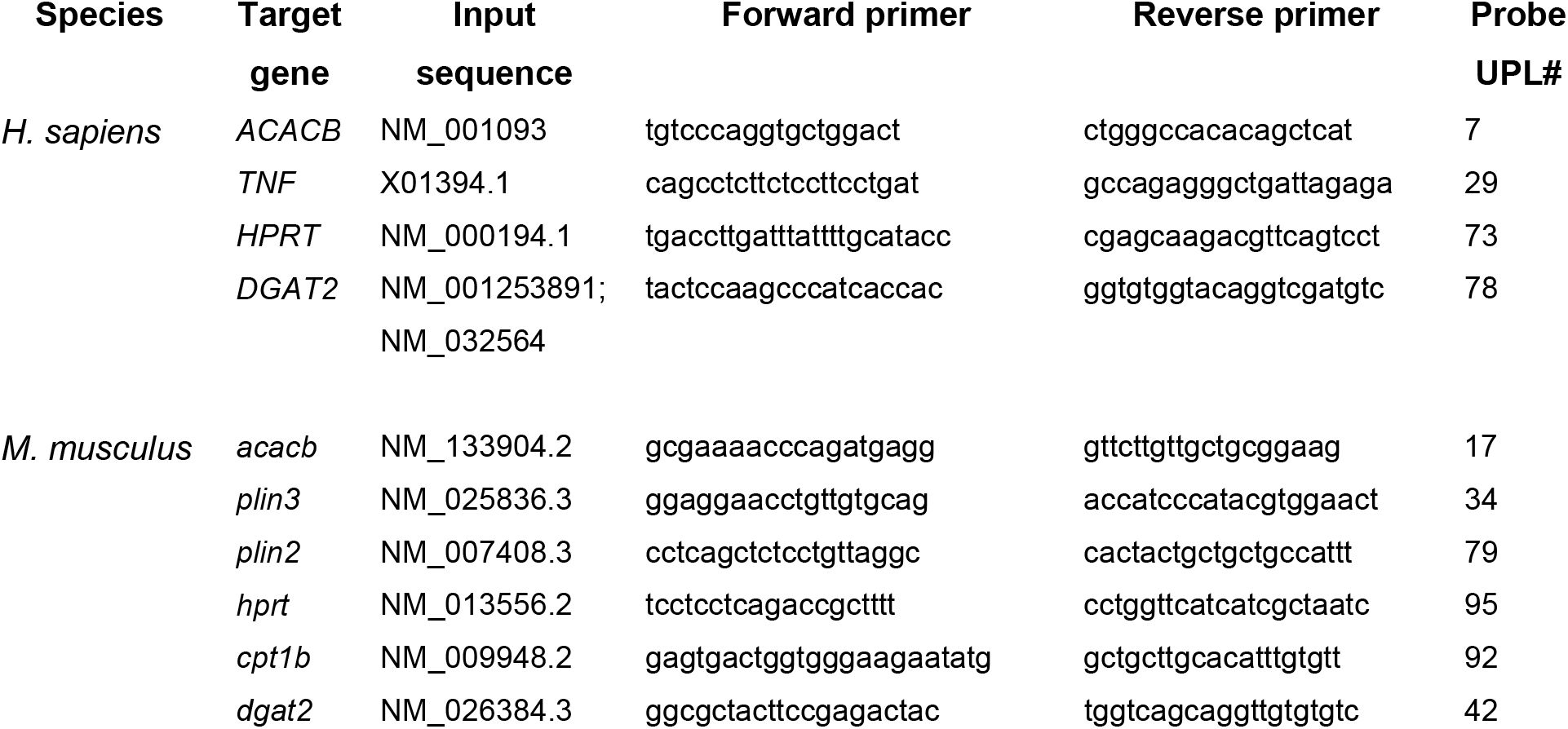
Primers used for qRT-PCR

### Microarray analyses

Integrity of extracted, total RNA was analyzed with the RNA Nano 6000 Kit on a Bioanalyzer (Agilent Technologies, Santa Clara, CA, USA) according to manufacturer’s instructions. Total RNA was used for reverse amplification and Cy3-labelling of cRNA as well as hybridization on Agilent Mouse Whole Genome 4×44K V2 arrays and scanning was conducted as described elsewhere^95^. GeneSpring version 12.6 (Agilent Technologies) was used for analysis of data with removal of compromised probes prior to analysis. Differences in gene expression were computed using a Moderated t-test with a Benjamini-Hochberg multiple comparison correction cut-off of p≤0.05 between infected *Wnt6*^+/+^ and *Wnt6*^−/−^ macrophages. Gene Symbols of significantly regulated genes (data available on request) were used to query the Molecular Signatures Database v6.0 (http://software.broadinstitute.org/gsea/msigdb) for enrichment of Reactome gene sets with a FDR q-value cut-off of p≤0.05.

### Immunohistochemistry

Lung tissue from patients with a multi-drug resistant TB was surgically removed (University Hospital Schleswig-Holstein (UKSH), Lübeck, Germany), dissected and fixed with 10% formalin for 24-48 hours. For immunohistochemical stainings, paraffin-embedded tissue was cut in 1 μm sections on a microtome (SM 2000R, Leica Biosystems), and sections were mounted on glass slides (SuperFrost Plus, R. Langenbrink, Emmendingen, Germany). Following de-parrafinization and antigen retrieval, which was performed at 90°C for 30 minutes in the presence of 10 mM citric acid, endogenous peroxidase activity was quenched by incubation with 3% H2O2 for 10 minutes. Slides were incubated in Antibody Diluent (Zytomed Systems, Berlin, Germany) in the presence of a primary antibody specific for WNT6 (purchased from Abcam (ab50030, 5 μg/ml) or Bio-Techne (AF4109, 6.6 μg/ml)), CD68 (clone PG-M1, 1:100, purchased from Agilent Technologies), PLIN2 (Abcam (# ab78920, 1:100), ACC2 (LS-C11360; LSBio, Seattle, USA) and ACC1/2 (mAb, C83B10; Cell signalling, Frankfurt, Germany). If necessary, tissue slides were incubated in Antibody Diluent (Zytomed Systems) containing a specific secondary antibody (F(ab)2 Fragment Rabbit Anti-sheep (Jackson Immoresearch, Suffork, UK) or rabbit anti-mouse IgG (Zytomed Systems) both 1:500 in Antibody Diluent) for 30-60 minutes. For detection and visualization, a Horseradish-Peroxidase (HRP)-conjugated Polymer based detection system (ZytoChem-Plus Kit Anti-rabbit) and the chromogene 3-amino-9-ethylcarbazole (AEC) (both from Zytomed Systems) were used according to the manufacturer’s instructions. Frozen lung tissue sections (5 μm) were air-dried, fixed (10% [v/v] ice-cold formalin) and mounted on glass slides. Subsequently, tissue was incubated for 20 minutes in 20% oil red O solution (Sigma) after washing with 60% 2-propanol (Sigma) in order visualize lipid droplets by light microscopy. All slides were counterstained with Gills hematoxylin (Vector, Lörrach, Germany) and analyzed with a BX41 microscope (Olympus, Hamburg, Germany) and the NIS-Elements software (NIS-Elements D3.10, SP3; Nikon).

### Immunofluorescence and flow cytometry analysis

For immunofluorescence analyses, cells were seeded on Chamber Slides^®^ (Lab Tek II, 8 well, Thermo Fisher). To monitor acidification of bacteria-containing compartments, macrophages were infected with GFP-expressing *M. tuberculosis* (see above) for 2 hours, incubated with 400 nM LysoTracker^®^ dye (DND-99, Thermo Fisher) for 2 hours and were thoroughly washed with PBS. Subsequently, cells were fixed with 1% (w/v) Paraformaldehyde for 24 hours (4°C). To block unspecific protein binding sites and permeabilize cells, slides were incubated in PBS containing 10% normal serum (Pan-Biotek) and 0.2% Triton-X100 for 1 hour. Lipid droplets and nuclei were stained with 0.2% Triton-X100/PBS containing 4,4-difluoro-1,3,5,7,8-pentamethyl-4-bora-3a,4a-diaza-s-indacene (BODIPY 493/503, 5 μg/ml; Thermo Fisher)^88^, and DAPI (1 μg/ml; Roche Applied Science), respectively, for 1 hour. For quantification of acidified, LysoTracker^®^ positive compartments, samples were evaluated in a blinded fashion (counting of >300 phagosomes per condition).

To compare neutral lipid content of cells by fluorescence microscopy, macrophages were infected with mCherry-expressing *M. tuberculosis*, fixed, and stained with BODIPY as described above. Subsequently, cells were visualized by fluorescence microscopy and obtained images were analyzed with ImageJ software (Version 1.51n) using a macro script (available on request). To assess relative changes in neutral lipid content, the area of the BODIPY signal was normalized to the nuclear area of the cells. For this purpose, nuclei were identified by DAPI staining and the nuclear area calculated for every image. The threshold to determine the BODIPY positive area within the images was determined by measuring the background signal within the nuclear area, which was essentially devoid of neutral lipids. A BODIPY signal above the average background multiplied by two times the standard deviation was considered positive. At least 200 cells per condition in each individual experiment were analyzed.

To visualize WNT6 and neutral-lipids in frozen lung tissue, sections (5 μm) were air-dried, fixed (10% [v/v] ice-cold formalin) and mounted on glass slides. Subsequently, unspecific protein binding sites were blocked by incubating sections with PBS containing 10% normal donkey serum (Pan-Biotek), 2% BSA and 0.2% Triton-X100 for 1 hour. Slides were incubated sequentially with an antibody specific for WNT6 (Bio-Techne (AF4109, 6.6 μg/ml)) and a suitable fluorescence (Cy3)-labelled secondary antibody (AffiniPure Donkey Anti-Sheep IgG, Minimal Cross Reactions, Jackson Immunoresearch, Cambridge, UK) for 2 and 1 hour, respectively. Neutral lipids and nuclei were visualized by use of BODIPY (10 μg/ml) and DAPI (1 μg/ml) as described earlier in this section. All slides were mounted with ProLong^™^ Antifade Reagents (Thermo Fisher), covered with glass coverslips (R. Langenbrink) and analyzed by use of an Axio Observer microscope, equipped with an ApoTome, and the AxioVision Software 4.8 or earlier (Carl Zeiss, Oberkochen, Germany).

To quantify neutral lipids by flow cytometry, NIH3T3 cells were detached by incubation with Accutase (Thermo Fisher) for 5 minutes at 37°C. Subsequently, cells were stained with BODIPY (5 μg/ml) for 1 hour, washed, re-suspended in PBS containing 0.2% EDTA and subjected to a MACS Quant Analyzer 10 (Milteny Biotec, Bergisch Gladbach, Germany) using the Milteny MACSQuantify software (Version 2.6 or 2.8). Data was analyzed with FCS Express v6 or earlier (De Novo Software, Glendale, CA, USA).

To determine mitochondrial activity, transdifferentiated BLaER1 WT and ACC2 knockout cells or hMDMs were either left untreated or infected at an MOI of 0.1 for 3 days. After washing with PBS, cells were stained with the membrane potential-sensitive dye Rhodamine 123 (25 minutes, 0.5 μg/ml) and the membrane potential-independent dye MitoTracker® Deep Red FM (300 nM) to measure the mitochondrial activity and mitochondrial mass, respectively (both Thermo Fisher). Cell were washed once and immediately analyzed on the FACS Canto II (BD) using the BD Diva Software (Version 6.1.2.). Data was analyzed with FCS Express v6 or earlier (De Novo Software, Glendale, CA, USA).

### Sample preparation and lipid extraction

For mass spectrometry based quantification of lipids from *in vitro* cultures, NIH3T3 cells were detached and transferred into suitable tubes (SafeLock, Eppendorf, Hamburg, Germany) in aliquots of 0.3-0.6 × 10^6^ cells. Subsequently, cells were washed with PBS at 37 °C. In order to remove residual liquid, cells were centrifuged (10.000 × *g*), and dry pellets immediately stored at −80 °C. BMDM (0.5 × 10^6^) were incubated in the presence of fatty acids or respective controls, washed with PBS at 37 °C, detached on ice for 1 hour, and treated and stored as described earlier for NIH3T3 cells.

For shotgun lipidomics analysis of infected mouse lungs, homogenates (200 μl in PBS / Protease-Inhibitor cocktail (Protean, Roche) were transferred into suitable tubes, incubated in methanol (800 μl, 2 h, RT) and stored at −80 °C until lipid extraction.

Total lipids were extracted according to a customized methyl-tert-butyl ether (MTBE) method^96^. Briefly, samples were dried in a SpeedVac and solved in 20 μl of 50 mM ammonium acetate. Then 270 μl methanol, containing 3% acetic acid, were added. After vortexing the mixture, internal standard solution (either, SPLASH® Lipidomix® Mass Spec Standard (330707, Avanti Polar Lipids, Alabaster, US) or standard mixture according to Table II) were added. Afterwards, 1 ml of MTBE was added and the solution was incubated for 1 hour at room temperature with continuous shaking at 600 rpm (Eppendorf, MixMate). Next, 500 μl of water was added and subsequently incubated for 10 mins at room temperature with continuous shaking at 1300 rpm. For phase separation, were centrifuged for 10 min at 15.000 × *g* and then the upper phase was collected in a separate tube. The lower phase was re-extracted with 400 μl theoretical upper phase, vortexed and incubated for 20 min at room temperature with continuous shaking (1300 rpm). The solution was once more centrifuged as described above. The resulting upper phases were combined and subsequently dried in a SpeedVac (Thermo Fisher Scientific, Waltham, US). The dried extracts were dissolved in a mixture of chloroform, methanol and water (60/30/4.5; v/v/v) and stored at −80 °C.

**Table II.**
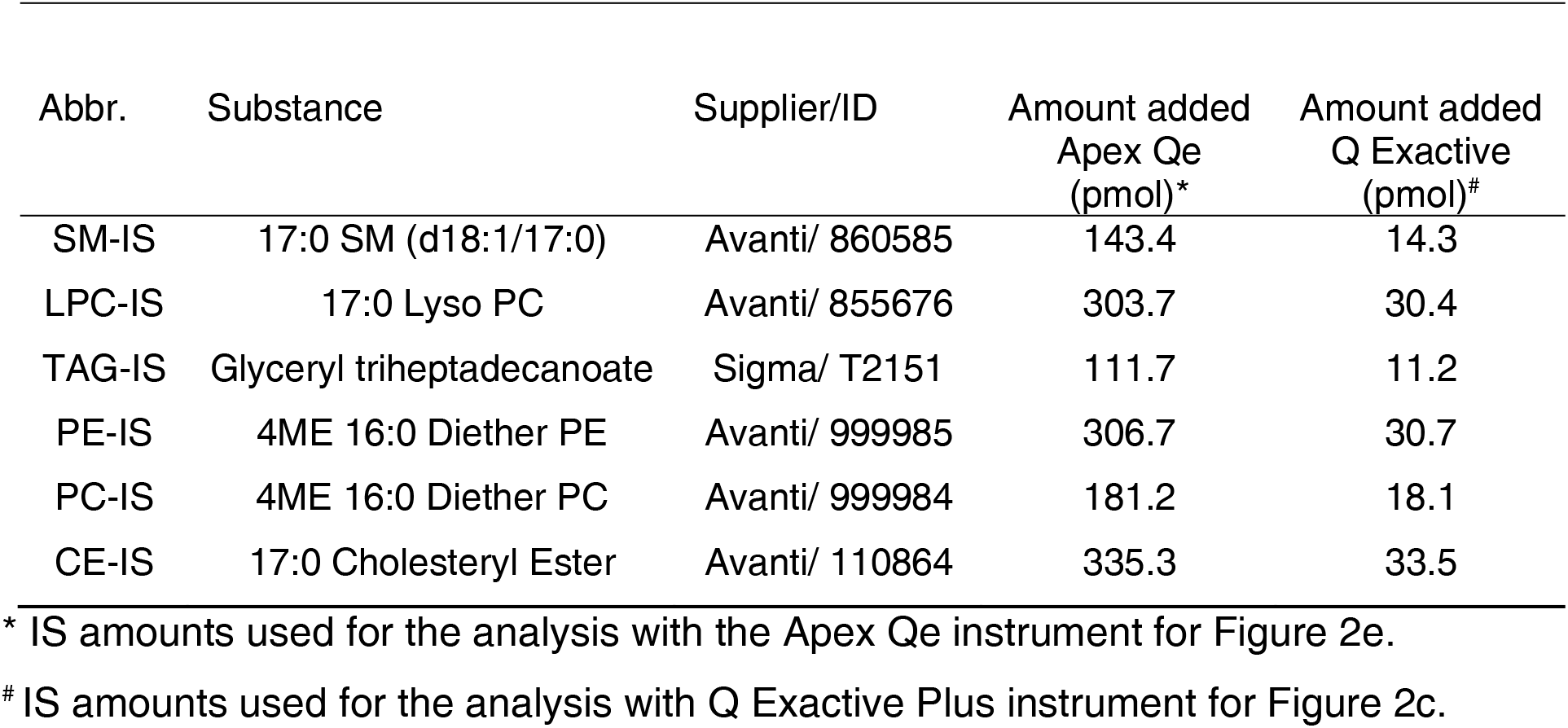
Internal standards used for lipid quantification of *in vitro* cultivated cells.

### Lipidomics

Shotgun lipidomics measurements were performed using a Q Exactive (Thermo Fisher Scientific, Bremen, Germany) or an Apex Qe Fourier Transform Ion Cyclotron Resonance mass spectrometer (Bruker Daltonik, Bremen, Germany), both equipped with a TriVersa NanoMate (Advion BioSciences, Ithaca, NY, USA) as autosampler and ion source^96,97^. Lipid identification was performed using LipidXplorer^98^ and quantitation was achieved in reference to a mix of internal standards, which were added prior extraction (Table II, SPLASH® Lipidomix® Mass Spec Standard).

### Tracing experiments with ^13^C-oleic acid

Uniformly ^13^C-labelled oleic acid (U-13C18, 98%, Cambridge Isotope Laboratories, Tewksbury, USA) was solubilized in ethanol (pure, for molecular biology, AppliChem) and conjugated to BSA (^13^C-oleate-BSA) as described earlier in this section. Cells were pulsed with ^13^C-oleate-BSA during differentiation of monocytes into macrophages (see protocol above). Subsequently, hMDMs were cultivated in the absence of isotope-labelled substrates and infected with Mtb for 4 hours. After removing extracellular bacteria by washing, macrophages were incubated for a total of 7 days in the absence or presence of ACC2 inhibitor 3 (ND-646). Finally, cells were detached on ice, washed and lysed by incubating in methanol (≥99% Chromasolv™) for 2 hours at RT. Lipids were extracted and quantified using shotgun lipidomics as described earlier. Briefly, ^13^C-labeled oleic acid incorporation in macrophages was traced by high resolution MS^1^ using the Q Exactive Plus. Incorporation rates for hMDMs were determined for the lipid classes PC, PC-O, TAG, CE and SM using the positive ion mode (Supplement Figure I). Quantitation was performed in reference to SPLASH® Lipidomix®. Metabolization of ^13^C-labeled OA in Mtb was traced using the major abundant phospholipid PI 16:0_19:0 (TSA) with a semi-targeted lipid analysis in the negative ion mode. The isotopic labelled 12C_1_13C_18_ TSA fragment in MS^2^ (*m/z* 315.34) and the ^12^C19 signal (*m/z* 297.28) were utilized to determine incorporation rates (preprint: Heyckendorf et al. Biorxiv, 2020).

### Cell viability assay

Real-time impedance measurements were conducted on a xCELLigence System (ACEA Bioscience, San Diego, USA) using plates with incorporated sensor array (E-Plate) and the Real-Time Cell Analyzer SP instrument. Data obtained were analyzed using the Real-Time Cell Analyzer Software 1.2 (ACEA Bioscience).

### Nitrite and cytokine quantification

To determine the production of reactive nitrogen intermediates (RNI), supernatants of *in vitro* cultivated cells were harvested and the content of nitrite was determined after adding Griess reagents by photometric measurement (absorbance at 540nm) on a Synergy2 (Biotek) microplate reader as described previously^99^.

To determine cytokine levels in Mtb-infected mouse lungs, homogenates were analyzed with a bead-based assay panel (Mouse Pro-inflammatory chemokine and mouse Inflammation Panel (LEGENDplex™), BioLegend, San Fransisco, USA) according to the manufacturer’s instructions. Measurements were performed on a FACSCanto™II (BD) flow cytometer and data were analyzed using the FCAP Array™ Software Version 3.0 (BD).

### Extracellular flux analysis

1.5×10^5^ BMDM were seeded on XF24 cell culture plates and incubated for 24h in the presence of cell culture medium containing BSA or Oleic acid (200 μM), which was complexed to BSA as described above. After washing and incubation with unbuffered DMEM containing 25 mM D-Glucose (Carl Roth, Karlsruhe, Germany) and 1 mM Pyruvate (Merck) for 1h at 37°C, cells were subjected to a XF24 Seahorse Analyzer (Agilent Technologies). During measurements Oligomycin (1 μM), FCCP [carbonyl cyanide 4-(trifluoromethoxy) phenylhydrazone] (1.5 μM) and Rotenone/Antimycin A (1 μM) (purchased from Agilent Technologies) were injected. Obtained data was analyzed by use of the Seahorse XF24 Software V 1.8.1.1.

### Ethics

All experiments performed with primary human cells or human lung tissue were reviewed and approved by the Ethics Committee of the University of Lübeck, Germany (#14-032,#12-220,#14-225,#18-194). All animal experiments were performed according to the German animal protection laws and were approved by the Animal Research Ethics Board of the Ministry of Environment (Kiel, Germany).

### Statistical analysis

Statistical analyses were performed using GraphPad Prism 7 or earlier software versions (GraphPad Software, La Jolla, CA). For statistical analyses of *in vitro* experiments, data was log-transformed in order to assume parametric distribution^100^. For group comparison, a Repeated Measure One-way ANOVA followed by Holm-Sidak multiple comparison as post-hoc test was performed. For statistical analysis of *in vivo* experiments, data was tested for normality, log-transformed and analyzed by an unpaired, one-tailed^101^ Student’s t-test. *p,0.05, **p, 0.01, ***p, 0.001. All data are shown as mean +/− SEM.

